# Ferroptotic vulnerability of post-meiotic germ cells skews sex chromosome ratios in aged testis

**DOI:** 10.1101/2024.10.21.619554

**Authors:** Jasper K. Germeraad, Takako Kikkawa, Junya Ito, Leon S.Y. Giesselink, Wilfred T.V. Germeraad, Bernhard Henkelmann, Maceler Aldrovandi, Kenshiro Hara, Kentaro Tanemura, Kiyotaka Nakagawa, Eikan Mishima, Marcus Conrad, Noriko Osumi

## Abstract

Age-related testicular germ cell depletion has been predominantly attributed to apoptosis; however, the contribution of alternative cell death modalities remains unclear. Using young and aged mice, we demonstrate compartment-specific cell-death signatures in the testis. Apoptotic cell increased in peripheral germ layers, whereas the luminal round spermatid (RS) layer mainly accumulated non-apoptotic events sharing features with ferroptosis, a lipid-peroxidation dependent cell death. Lipidomic analysis revealed age-related phospholipid remodelling in whole testis, while RS-enriched fractions showed increased lipid peroxidation. Interestingly, aging shifted the balance of sex-chromosome–bearing RS toward Y-bearing cells, and this bias persisted in mature sperm. Dietary vitamin E modulated these phenotypes bidirectionally: vitamin E deficiency in young mice increased RS lipid peroxidation and reproduced the Y-skew, whereas supplementation in aged mice lowered RS lipid peroxidation, decreased non-apoptotic events, and restored the skewed RS sex-chromosome ratio. These findings define an age-related RS vulnerability linking redox stress to haploid spermatogenic output.

## Introduction

Advanced paternal age is associated with reduced semen volume, lower sperm concentration and motility, and increased sperm DNA fragmentation^1–3^. Although male fertility can be maintained throughout life, reproductive capacity progressively declines because of age-dependent deterioration of testicular function^4^. Sperm are produced through spermatogenesis, a strictly controlled process within the seminiferous tubules in which spermatogonial stem cells generate differentiating spermatogonia, meiotic spermatocytes and post-meiotic haploid round spermatids (RSs) that subsequently elongate into spermatozoa^5^. Each round spermatid (RS) carries either a Y- or an X-chromosome. Therefore, selective survival or loss at post meiotic stages has the potential to bias sex-chromosome transmission by altering the relative representation of Y- and X-bearing gametes. While age-related impairment of murine spermatogenesis is well documented^6–8^, the mechanisms driving germ-cell loss during testicular aging remain insufficiently understood, particularly at post-meiotic stages^9^.

Germ-cell loss in aging testes has frequently been linked to oxidative stress-induced apoptosis^10,11^. However, oxidative stress can activate multiple cell death modalities^12^, and susceptibility is likely to vary along the germ-cell differentiation axis. Previous reports showed that pre-meiotic germ and somatic cells predominantly engage apoptosis^13^, whereas recent *in vitro* studies suggests that spermatocytes and RSs respond differently to oxidative insults, including lipid peroxides^14^. RS membranes are enriched in polyunsaturated phospholipids and undergo extensive membrane remodelling, features predicted to increase sensitivity to lipid peroxidation^15,16^. Consistent with this notion, elevated lipid peroxidation represents a recurring biochemical signature of reproductive aging in testes^17^.

Lipid peroxidation can trigger ferroptosis, a non-apoptotic form of cell death driven by iron-mediated phospholipid peroxidation^18^. Ferroptosis susceptibility is shaped by three core pillars: antioxidant defence mechanisms, membrane phospholipid composition and control of labile iron^19,20^. For instance, the selenoenzyme glutathione peroxidase 4 (GPX4) detoxifies potentially harmful lipid hydroperoxides, and impaired GPX4 functions confers cellular vulnerability to ferroptosis^21,22^. Conversely, long-chain fatty acid CoA ligase 4 (ACSL4) promotes incorporation of poly-unsaturated fatty acids (PUFAs) into membrane phospholipids, generating highly peroxidizable substrates that sensitize cells to ferroptosis^23^. Iron-handling pathways further modify this process; transferrin receptor 1 (TFR1) mediated iron uptake and heme oxygenase-1 (HO-1) dependent heme degradation can elevate iron levels, amplifying ferroptotic sensitivity^24^.

Ferroptosis is suppressed by lipophilic antioxidants, among which vitamin E is one of the most potent natural antioxidants that limits membrane lipid peroxidation and exerts anti-ferroptotic activity^25^. Vitamin E deficiency induces spermatogenic defects resembling features of reproductive aging^26^ and can preferentially reduce post meiotic germ-cell populations, despite differences in dietary regimens and species^27^. These observations suggest that disruption of lipid redox homeostasis may be sufficient to activate aging-relevant vulnerability pathways, with RSs representing a particularly susceptible stage where lipid peroxidation could selectively influence the survival of X-versus Y-bearing haploid cells.

Here, we investigate how aging alters spermatogenic output and cell death modalities, including apoptosis and ferroptosis, across the seminiferous epithelium. We characterize age-related lipid peroxidation and related defence pathways in whole testis and in RS-enriched fractions and quantify sex-chromosome ratios in both RSs and mature sperm. Additionally, by bidirectionally manipulating dietary vitamin E, through deficiency in young mice and supplementation in aged mice, we test whether aging-associated RS vulnerability and sex-chromosome ratio shifts are causally linked to the dysregulation of the lipid peroxidation-ferroptosis axis.

## Results

### Aging impairs spermatogenesis and promotes non-apoptotic cell death in RSs

By 18 months of age, C57BL/6J males exhibit measurable testicular aging^28^. To investigate aging-related gonadal changes, we therefore compared young (3-month-old) and aged (18-month-old) male mice. Aged mice displayed increased body weight (Fig. 1a) and decreased testis weight (Fig. 1b), resulting in a lower gonadosomatic index (testis weight/body weight, Fig. 1c). To assess spermatogenic output in aged mice, we focused on stage VII-VIII seminiferous tubules, which contain undifferentiated spermatogonia, differentiating spermatogonia, meiotic pachytene spermatocytes, post-meiotic RSs and somatic Sertoli cells (spermatogenesis support cells), thus representing key stages of spermatogenesis (Fig. 1d). Testis from aged mice showed reduced cellular density in germ and somatic cells of the seminiferous epithelium (Fig. 1e). Quantitative analysis showed a reduction in undifferentiated spermatogonia in aged compared with young testes (Fig. 1f), which extended to differentiating spermatogonia, pachytene spermatocytes, RSs, and was also observed in somatic SCs (Fig. 1g–j).

**Fig. 1:**
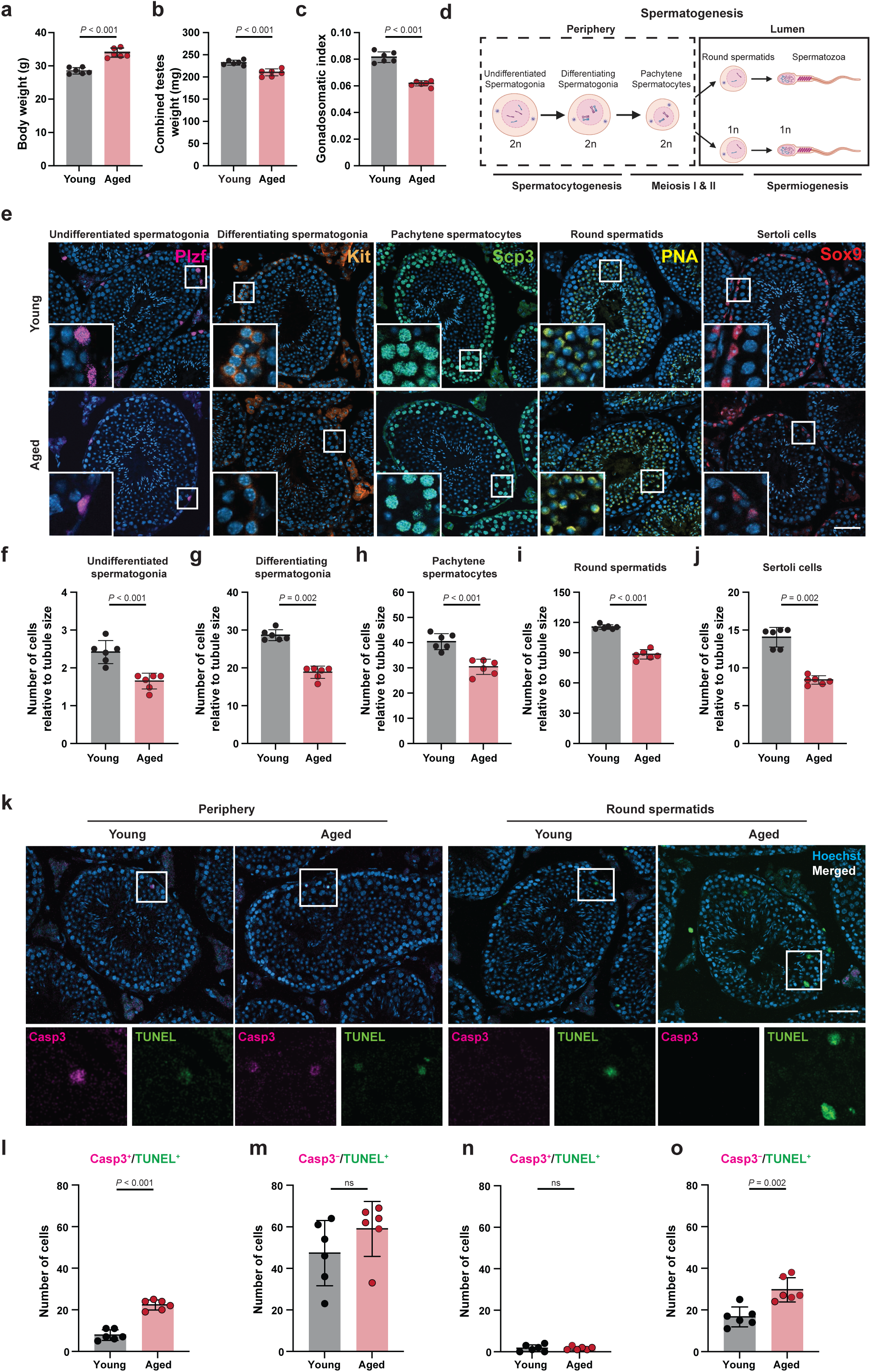
Age-related alterations in mouse testis germ cell populations. **a**–**c**, Body weight (**a**), combined testes weight (**b**) and gonadosomatic index (combined testes weight/body weight; **c**) in young (3 months) and aged (18 months) males. **d**, Schematic of spermatogenesis highlighting undifferentiated and differentiating spermatogonia, pachytene spermatocytes, round spermatids (RSs) and spermatozoa. **e**, Representative stage VII–VIII seminiferous tubules stained for PLZF (undifferentiated spermatogonia), KIT (differentiating spermatogonia), SCP3 (pachytene spermatocytes), PNA (RSs) and SOX9 (Sertoli cells); nuclei counterstained with DAPI. Scale bar, 50 µm. f–j, Quantification of PLZF^+^ (**f**), KIT^+^ (**g**), SCP3^+^ (**h**), PNA^+^ (**i**) and SOX9^+^ (**j**) cells, normalized to tubule circumference (**f**: 20 images per mouse; **g**–**j**: 10 images per mouse). **k**, Representative tubules showing CASP3^+^/TUNEL^+^ and CASP3^−^/TUNEL^+^ events in peripheral and luminal compartments; nuclei counterstained with Hoechst. Scale bar, 50 µm. **l**–**o**, Quantification of CASP3^+^/TUNEL^+^ and CASP3^−^/TUNEL^+^ cells in the periphery (**l**, **m**) and lumen (**n**, **o**), normalized to 100 tubules. Data are mean ± s.d.; dots represent mice. Statistics: two-tailed Student’s *t*-test (**a**–**c**, **f**, **h**, **i**, **l**, **n**, **o**) or Mann–Whitney *U* test (**g**, **j**, **m**), exact *P* values are given. *N* = 6 mice per group. Illustration in **d** was created in BioRender, Germeraad, J. (2026) https://BioRender.com/8iim1z1

Next, to investigate germ cell type–specific cell death associated with aging, we evaluated cell death using terminal deoxynucleotidyl transferase dUTP nick end labelling (TUNEL) across seminiferous tubules, irrespective of stage. Although TUNEL staining was classically used as a marker of apoptosis, it is also known to detect cells undergoing other regulated cell death modalities^19^. A higher proportion of seminiferous tubules in aged mice contained TUNEL-positive dead cells compared to those in young mice (Extended data Fig. 1a, b).

To identify apoptotic cells, we performed double labelling with TUNEL (a general cell death marker) and immunostaining against cleaved-caspase3 (CASP3), a specific marker of apoptosis. We quantified CASP3^+^/TUNEL^+^ cells (apoptotic) and CASP3^−^/TUNEL^+^ cells (dead cells) separately within the peripheral compartment, containing spermatogonia, spermatocytes and Sertoli cells, and the luminal layer, which is enriched in RSs (Fig. 1k). In the peripheral compartment, aging mice showed a significantly larger number of CASP3^+^/TUNEL^+^ apoptotic cells (Fig. 1l), whereas the number of CASP3^−^/TUNEL^+^ dead cells remained unchanged (Fig. 1m). In contrast, within the luminal layer, CASP3^+^/TUNEL^+^ apoptotic RSs were rare and did not differ between age groups (Fig. 1n, Extended Data Fig. 1c), while CASP3^−^/TUNEL^+^ cells prevailed and were more abundant in aged testes (Fig. 1o). Although cleaved CASP3 can be transient, the marked enrichment of CASP3^−^/TUNEL^+^ events specifically in the luminal RS layer supports a compartment-specific shift toward TUNEL^+^ cell-death events without detectable CASP3 activation in aging testes.

### Aging enhances lipid peroxidation in RSs

Increased lipid peroxidation is a feature of testicular aging and capable of inducing cell death in aging rodent models^29–31^. To define age-related changes in phospholipid composition and oxidation status, we profiled testicular phospholipids by mass-spectrometry-based lipidomic analysis. Consistent with previous studies^32,33^, phosphatidylcholine (PC) and phosphatidylethanolamine (PE) constituted the predominant phospholipid classes detected in mouse testis (Extended Data Fig. 2a, b). Age-related changes in individual PC and PE species were modest overall, although aged testes showed an increase in PC plasmalogen species (Extended Data Fig. 2c), but not PE plasmalogen species (Extended Data Fig. 2d). The increase in PC plasmalogen species may reflect age-dependent remodelling of ether lipids, which are implicated in antioxidant defence under oxidative stress conditions^34^.

Given that RSs are enriched in polyunsaturated phospholipid species^15^, this may suggest heightened vulnerability to oxidative stress and lipid peroxidation. We isolated RS-enriched fractions from young and aged testes using a density-gradient (Fig. 2a) and confirmed >80% RS purity by peanut agglutinin (PNA) staining and nuclear morphology (Extended Data Fig. 3). Immunoblot analysis of RS-enriched fractions revealed increased levels of 4-hydroxynonenal (4HNE)-modified proteins in aged compared with young samples, indicating enhanced lipid peroxidation in aged RSs (Fig. 2b). In line, gene expression levels of *Alox15*, encoding a PUFA-oxidizing lipoxygenase, was elevated in aged RSs (Fig. 2c). Because susceptibility to lipid peroxidation is influenced by how PUFAs are incorporated into membrane phospholipids^23^, we examined gene expression levels of key enzymes involving PUFA loading and elongation. In aged RSs, expression of *Acsl4,* encoding an acyl-CoA synthetase that facilitates arachidonic/adrenic acid (AA/AdA)-PUFA enrichment in phospholipids, was increased, whereas other ACSL family genes such as *Acsl6* and the PUFA-elongating enzymes *Elovl2* and *Elovl5* were not altered with age (Fig. 2d, e, Extended Data Fig. 2e, f). Together, these data are consistent with a selective shift toward AA/AdA-type PUFA phospholipid remodelling rather than enhanced docosahexaenoic acid (DHA) loading in aged RSs, supporting increased susceptibility of AA/AdA-containing phospholipids to oxidation.

**Fig. 2:**
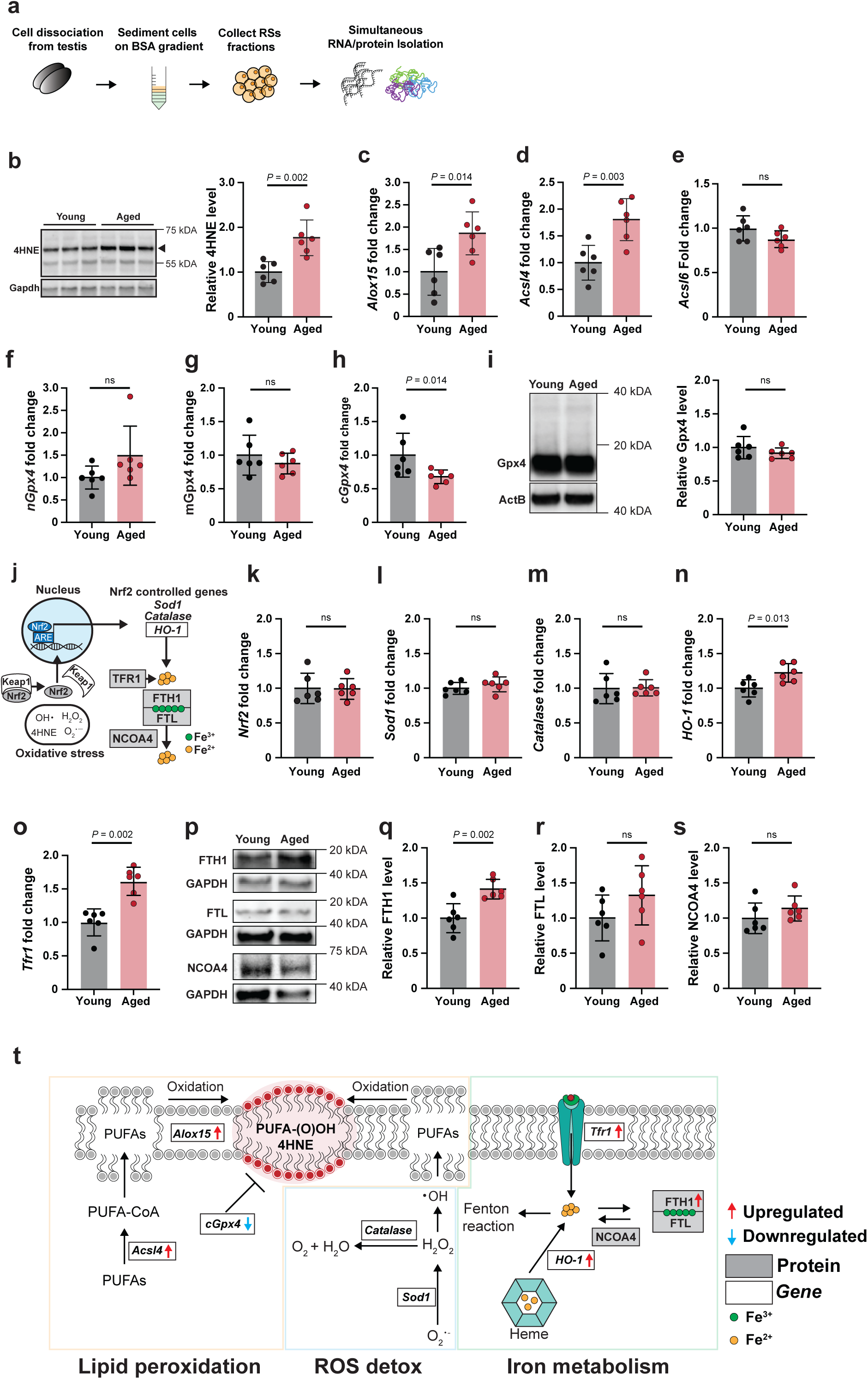
Increased lipid peroxidation in aged whole testis and round spermatids. **a**, Workflow for isolation of RS-enriched fractions and extraction of RNA/protein using a modified density-gradient approach. **b**, Representative immunoblots of 4HNE adducts and quantification, normalized to GAPDH, black triangle indicates quantified band. **c**–**h**, RS mRNA expression of *Alox15* (**c**), *Acsl4* (**d**), *Acsl6* (**e**) and *Gpx4* isoforms (*nGpx4* (**f**), *mGpx4* (**g**), *cGpx4* (**h**)), shown as fold change relative to *ActB*. **i**, Representative immunoblots and quantification of GPX4 protein, normalized to ACTB. **j**, Schematic outlining NRF2 activation and initiating its target gene expression. Oxidative stress activates NRF2, via KEAP1 release, to induce antioxidant/HO-1 programs, while iron uptake (TFR1), storage (FTH1/FTL), and ferritinophagy (NCOA4) regulate the labile iron pool (LIP) **k**–**o**, RS mRNA expression of *Nrf2* (**k**), *Sod1* (**l**), *Cat* (**m**), *HO-1* (**n**) and *Tfr1* (**o**), relative to *ActB*. **p**–**s**, Representative immunoblots (**p**) and quantification of FTH1 (**q**), FTL (**r**) and NCOA4 (**s**). **t**, Summary schematic of aging-associated changes across lipid peroxidation, redox defence and iron metabolism (red, increased; blue, decreased; gray boxes, protein; white boxes, mRNA). Data are mean ± s.d.; dots represent mice. Statistics: Welch’s *t*-test (**a**); two-tailed Student’s *t*-test (**b**–**e**, **g**–**i**, **k**, **l**, **n**, **q**–**s**); Mann–Whitney *U* test (**f**, **m**, **o**), exact P values are given. *N* = 6 mice per group.

### Aging reduces cGpx4 expression and remodels iron handling in RSs

To investigate the mechanism underlying increased lipid peroxidation in aged testes, we assessed antioxidant defence and iron-handling pathways in RSs. GPX4 is a central regulator of lipid peroxide detoxification and the prime regulator of ferroptosis. Immunostaining confirmed GPX4 expression in the RS cytoplasm and in the sperm tail (Extended Data Fig. 4a). In testis, GPX4 is expressed as three isoforms: nuclear (nGPX4), mitochondrial (mGPX4) and cytosolic (cGPX4) isoforms, of which cGPX4 has been implicated as the critical isoform protecting germ cell from cell death^35–38^. We therefore quantified *Gpx4* mRNA transcripts in RSs from young and aged mice, using primer sets targeting the nuclear (*nGpx4*) transcript and the mitochondrial-targeted (*mGpx4*) transcript, together with a primer set amplifying both mitochondrial and cytosolic (*cGpx4*) transcripts^39^. Using this strategy, we estimated the *cGpx4* fraction as the difference between the shared signal and the mitochondrial-specific signal. While *nGpx4* and *mGpx4* were unchanged with age, *cGpx4* expression was reduced in aged RSs (Fig. 2f–h). Total GPX4 protein levels, detected with an antibody recognizing all isoforms, did not differ between young and aged RSs fractions (Fig. 2i). Given the predominance of mGPX4 in RSs^36^, these findings suggest that age-associated downregulation of *cGpx4* occurs without a detectable change in bulk RS GPX4 protein levels.

Nuclear erythroid factor 2 (NRF2) controls a broad antioxidant and cytoprotective program^40^ (Fig. 2j). Among NRF2 target genes, superoxide dismutase-*1* (*Sod1*) and catalase (*Cat*) mRNAs did not change in RSs (Fig. 2k–m), whereas *HO-1* was selectively induced in aged RSs (Fig. 2n), consistent with increased heme catabolism. Because HO-1 can elevate the labile iron pool, we assessed iron handling and found increased *Tfr1* mRNA and increased ferritin heavy chain (FTH1), with no change in ferritin light chain (FTL) and nuclear receptor co-activator 4 (NCOA4) protein in aged RSs (Fig. 2o–s; Extended Data Fig. 4b, c). Together, these changes support an age-associated shift toward a more pro-oxidant iron environment in RSs (Fig. 2t), which would be expected to sensitize these cells to iron-dependent lipid damage.

### Aging skews the Y/X sex chromosome ratio in RSs and mature sperm

Since *Acsl4* is encoded by the X-chromosome and is upregulated in aged RSs (Fig. 2f), we asked whether aging is associated with a bias in the relative abundance of X- and Y-bearing RSs. To distinguish X- and Y-bearing RSs, we performed fluorescent *in situ* hybridization (FISH), targeting the X- and Y-chromosome with specific probes. We found that the ∼1:1 ratio was maintained in young testes but was significantly skewed toward Y-bearing RSs in aged testes (young: mean Y/X ratio 1.02 [*n* = 6,282±100 Y-RSs, *n* = *6*,188±113 X-RSs]; aged: mean Y/X ratio 1.11 [*n* = 6,694±85 Y-RSs, *n* = 6,056±98 X-RSs]; Fig. 3a,b). To determine whether this bias persisted in mature sperm, we quantified X- and Y-bearing sperm using fluorescent activated cell sorting (FACS) using Hoechst-33342 to visualize the DNA content difference between Y- and X-bearing RSs (Fig. 3c, d). Consistent with the findings in RSs, the Y/X ratio was similarly shifted toward Y-bearing sperm in aged males compared with young controls (young: mean Y/X 1.02 [*n* = 3,382,147 ± 140,662 Y-sperm vs *n* = 3,293,224 ± 126,522 X-sperm]; aged: mean Y/X 1.09 [*n* = 3,294,779 ± 156,878 Y-sperm; *n* = 3,032,412 ± 145,748 X-sperm], Fig. 3e,f). These results indicate that aging is associated with a bias toward Y-bearing RSs, and this sex chromosome bias is maintained in mature sperm.

**Fig. 3:**
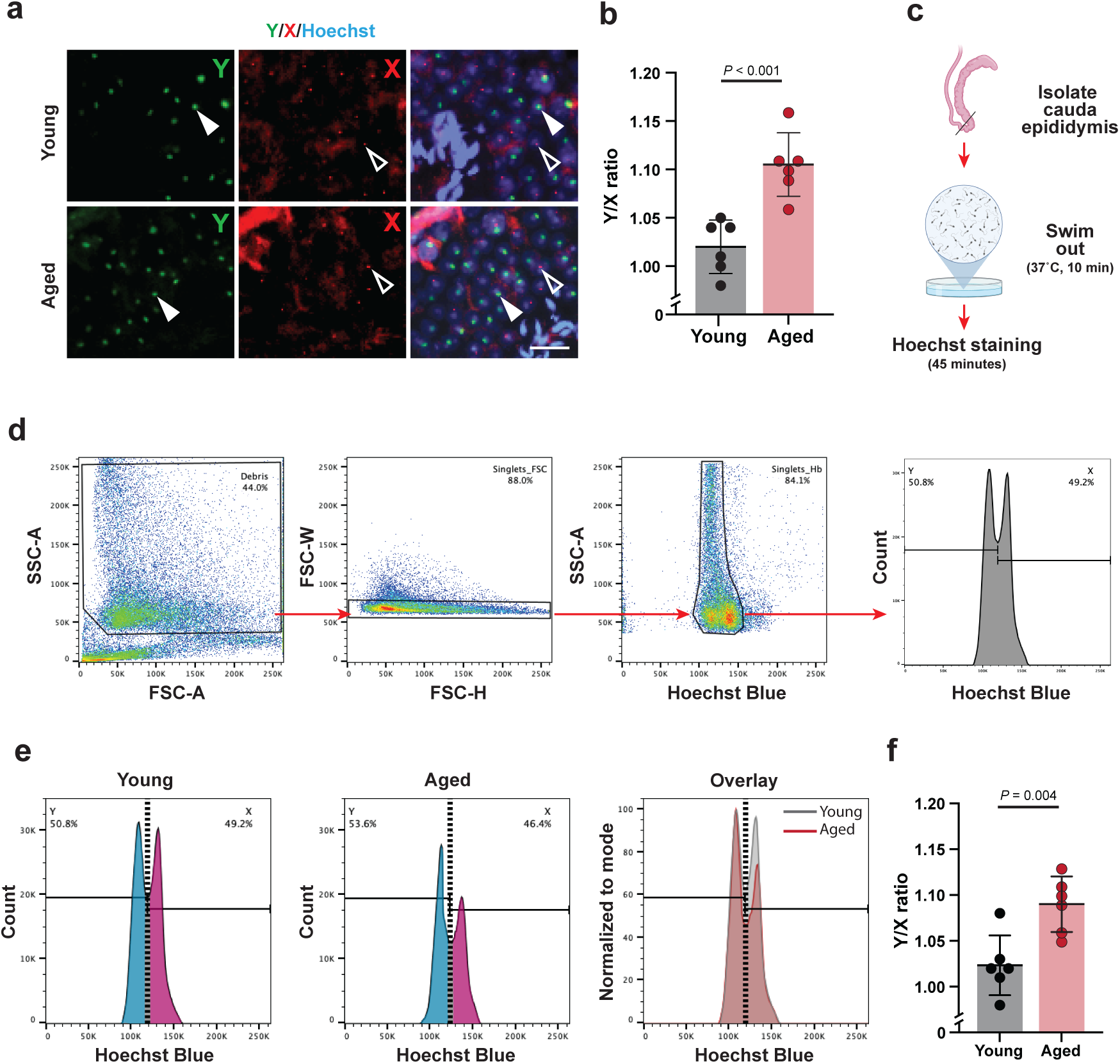
Aging alters the sex chromosome ratio in round spermatids and mature sperm. **a**, **b**, FISH in RSs showing Y-chromosomes (green; closed arrowhead) and X-chromosomes (red; open arrowhead) signals with Hoechst counterstain (**a**) and quantification of Y/X ratio (Young: n = 6,282±100 Y-RSs, *n* = *6*,188±113 X-RSs; mean Y/X ratio 1.02; Aged: *n* = 6,694±85 Y-RSs, *n* = 6,056±98 X-RSs; mean Y/X ratio 1.11) (**b**). Scale bar, 5 µm. **c**, Sperm collection and DNA staining workflow: cauda epididymides were excised and sperm was collected via swim-out (37 °C, 10 min) followed by Hoechst 33342 staining (45 min) before flow cytometry. **d**, Gating strategy to resolve X- and Y-bearing sperm: debris exclusion (FSC-A vs SSC-A), singlet selection (FSC-H vs FSC-W and Hoechst Blue vs SSC-A), and histogram-based separation by Hoechst Blue intensity. **e**, **f**, Representative Hoechst Blue distributions (young vs aged and overlay normalized to mode) (**e**) and Y/X ratio quantification (young: *n* = 3,382,147 ± 140,662 Y-sperm vs *n* = 3,293,224 ± 126,522 X-sperm; mean Y/X 1.02; aged: *n* = 3,294,779 ± 156,878 Y-sperm; *n* = 3,032,412 ± 145,748 X-sperm; mean Y/X 1.09) (**f**). Data are mean ± s.d.; dots represent mice. Statistics: two-tailed Student’s t-test (**b**, **f**). Exact *P* values are shown. *N* = 6 mice per group. Illustration in **c** was created in BioRender, Germeraad, J. (2026) https://BioRender.com/j1osdbj

### Vitamin E depleted diet recapitulated aging-associated phenotypes

Given the elevated lipid peroxidation observed in aged testis, we asked whether a lipid peroxidation-prone condition induced by vitamin E-depleted diet may cause aging-like testicular phenotypes in young mice^26^. Young animals were fed either a vitamin E-deficient diet (VE(−), <0.1mg/kg vitamin E) or control diet (135mg/kg vitamin E) for two cycles of spermatogenesis^5^ (70 days, Fig. 4a). Although body weight, testis weight and gonadosomatic index were not affected by vitamin E-deficient diet (Fig. 4b–d), VE(−) testes showed impaired spermatogenic output (Fig. 4e). Quantitative analysis revealed that numbers of undifferentiated and differentiating spermatogonia remained unchanged, whereas pachytene spermatocytes, RSs and Sertoli cells were reduced in VE(−) testes compared with controls (Fig. 4f–j), indicating impaired spermatogenic progression.

**Fig. 4:**
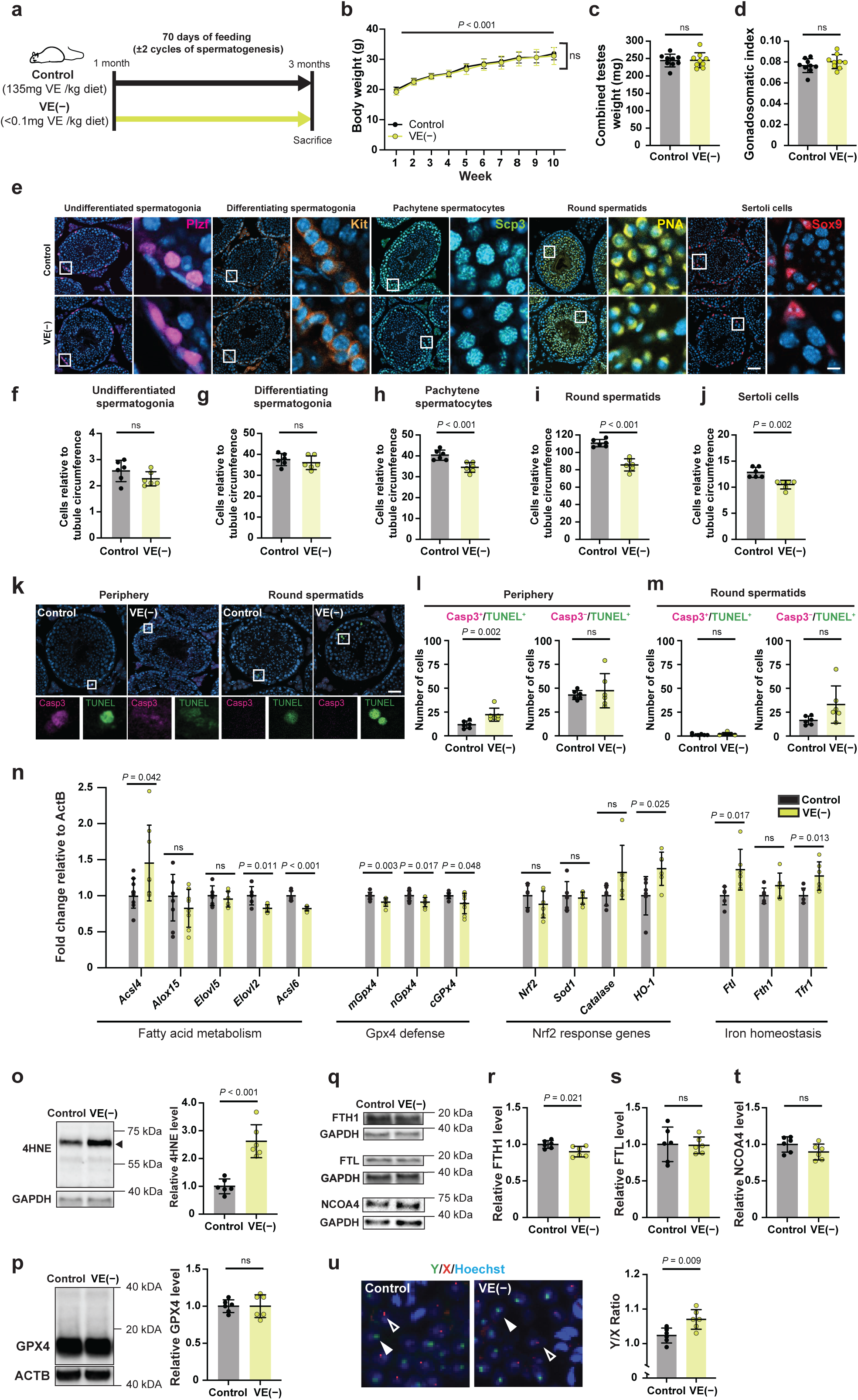
Vitamin E deficient feeding recapitulates age-associated testis phenotypes. **a**, Experimental design: young males (1 month) fed control or vitamin E–deficient diet (VE(−)) for 70 days. **b**, body weight was recorded weekly for 10 weeks in the same cohort of mice. **c, d,** combined testes weight (**c**) and gonadosomatic index (combined testes weight/body weight; **d**). **e**, Representative stage VII–VIII seminiferous tubules stained for PLZF, KIT, SCP3, PNA and SOX9: counterstained with DAPI. **f**–**j**, Quantification of PLZF^+^ (**f**), KIT^+^ (**g**), SCP3^+^ (**h**), PNA^+^ (**i**) and SOX9^+^ (**j**) cells, normalized to tubule circumference (**f**: 20 images per mouse; **g**–**j**: 10 images per mouse). **k**, Representative tubules showing CASP3^+^/TUNEL^+^ cells in the periphery and CASP3^−^/TUNEL^+^ cells in the lumen. **l**, **m**, Quantification of CASP3^+^/TUNEL^+^ and CASP3^−^/TUNEL^+^ cells in peripheral (**l**) and luminal compartments (**m**). **n**, qPCR of fatty-acid metabolism, GPX4 defence, NRF2-response and iron-homeostasis transcripts, relative to ActB. **o**, **p**, Representative immunoblots and quantification of 4HNE (**o**) and GPX4 (**p**), normalized to GAPDH, black triangle indicates quantified band. **q**–**t**, Representative immunoblots (**q**) and quantification of FTH1 (**r**), FTL (**s**) and NCOA4 (**t**). **u**, FISH in RSs with X (red) or Y (green) signals and Y/X ratio (control: *n* = 6,390±125 Y-RSs vs *n* = 6,274±127 X-RSs; mean Y/X ratio 1.02; VE(−): *n* = 6,548±222 Y-RSs vs *n* = 6,186±212 X-RSs; mean Y/X ratio 1.09). Scale bars: **e**, 50 µm (overview) and 5 µm (inset); **k**, 50 µm; **u**, 5 µm. Data are mean ± s.d.; dots represent mice except for **b** where it represents the mean of each group. Statistics: two-way repeated measures-ANOVA with Geisser–Greenhouse correction (**b**); two-tailed Student’s *t*-test (**c**, **d**, **f**–**i**, **m**–**p**, **r**–**t**, **u**); Mann–Whitney *U* test (**j**, **l**, **n**), exact *P* values are given. Sample sizes: **b**–**d**, *N* = 9 per group; all other panels, *N* = 6 per group.

Consistent with the reduction in germ and somatic cell numbers, VE(−) testes contained more TUNEL^+^ dead cells than control testes (Extended Data Fig. 5a, b). When cell death was analysed separately in the peripheral compartment and the luminal RS layer (Fig. 4k), vitamin E deficiency significantly increased CASP3^+^/TUNEL^+^ apoptotic cells in the peripheral regions without altering CASP3^−^/TUNEL^+^ cells (Fig. 4k, l). By contrast, in the luminal compartment, vitamin E-deficient diet only induced a trend toward higher CASP3^−^/TUNEL^+^ non-apoptotic RSs, but CASP3^+^/TUNEL^+^ RSs remained rare and unchanged (Fig. 4l, m; Extended Data Fig. 5d). Together, vitamin E deficiency preferentially elevates apoptosis-associated death events in the peripheral compartment, while its effect on RS-layer CASP3^−^/TUNEL^+^ events is weaker under these conditions.

In RSs isolated from VE(−) testes, expression of lipid-metabolic and stress-response genes was significantly altered. *Acsl4, Hmox1, Ftl* and *Tfr1* transcripts were upregulated, whereas all three *Gpx4* isoforms were reduced (Fig. 4n). Consistent with enhanced lipid peroxidation, the levels of 4HNE-modified proteins were elevated in VE(−) RSs (Fig. 4o). At the protein level, FTH1 was decreased, whereas total GPX4, FTL and NCOA4 were unaffected (Fig. 4p-t).

We further examined whether feeding mice with a vitamin E-deficient diet altered the sex chromosome composition of RSs. In control testes, Y- and X-bearing RSs were similar in number, whereas VE(−) testes exhibited a significant enrichment of Y-bearing RSs (control: mean Y/X ratio 1.02 [*n* = 6,390±125 Y-RSs vs *n* = 6,274±127], X-RSs; VE(−): Y/X ratio 1.07 [*n* = 6,548±222 Y-RSs vs *n* = 6,186±212 X-RSs]; Fig. 4u). Overall, vitamin E deficiency in young mice induced an aging-like RS stress state, with increased lipid peroxidation and broad remodelling of lipid/iron-defence transcripts in RS fractions. Notably, despite limited induction of RS-layer CASP3^−^/TUNEL^+^ events, vitamin E deficiency produced a significant shift in haploid output, enriching Y-bearing RSs, thereby recapitulating the sex-chromosome skew observed in aged testes.

### Vitamin E supplementation reduced aging-associated testicular phenotypes

Given that vitamin E deficiency in young mice recapitulated multiple aging-like testicular phenotypes, we next asked whether vitamin E supplementation could counteract these changes in aged mice. To this end, aged animals were fed either a vitamin E-supplemented diet (VE(+)) or control diet according to the regimen shown in Fig. 5a. Body weight, testis weight and gonadosomatic index did not differ between VE(+) and control groups (Fig. 5a–d), indicating that vitamin E supplementation does not affect gross somatic growth or testis size in aged mice during this feeding regimen. Despite unchanged testis size, VE(+) improved spermatogenic output in aged mice (Fig. 5e). While the numbers of undifferentiated and differentiating spermatogonia were not altered, pachytene spermatocyte, RS and Sertoli cell numbers were significantly increased in VE(+) compared with control testes (Fig. 5f–j). Consistent with this improvement, VE(+) testes contained fewer TUNEL^+^ cells than control testes (Extended Data Fig. 5a, c). In the peripheral compartment, vitamin E supplementation significantly lowered the Casp3^+^/TUNEL^+^ apoptotic population without affecting CASP3^−^/TUNEL^+^ cells (Fig. 5k, l). In contrast, within the luminal RS layer, VE(+) selectively reduced non-apoptotic CASP3^−^/TUNEL^+^ RSs, whereas CASP3^+^/TUNEL^+^ apoptotic RSs remained infrequent and unchanged across groups (Fig. 5k, m, Extended Data Fig. 5d). These results indicate that vitamin E attenuates apoptosis in the periphery and CASP3^−^/TUNEL^+^ non-apoptotic events in the luminal layer, consistent with reduced oxidative burden.

**Fig. 5:**
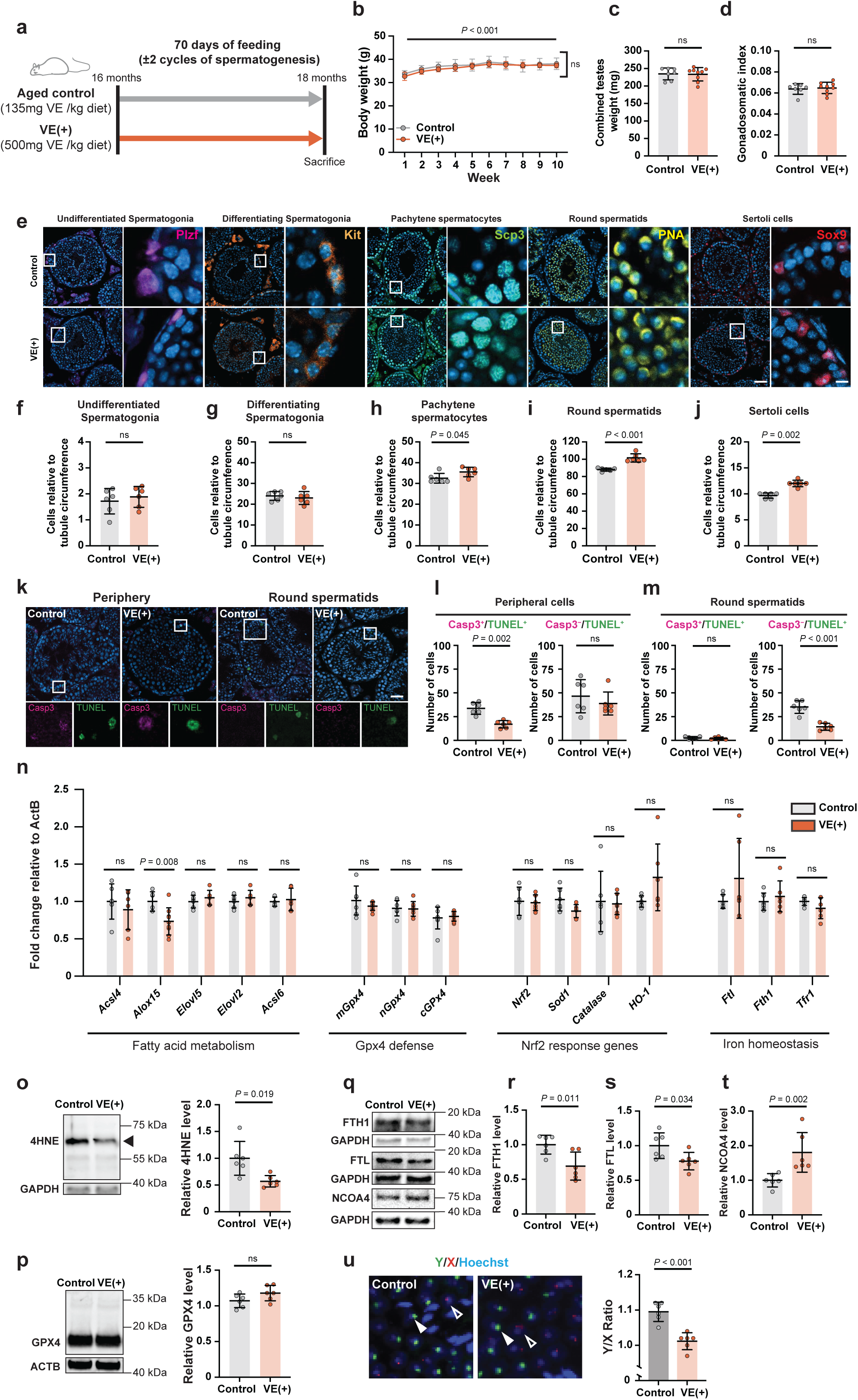
Vitamin E supplemented feeding rescues age-associated phenotypes within the testis. **a**, Experimental design: aged (16-months) males fed control or vitamin E–supplemented diet (VE+) for 70 days. **b**, body weight was recorded weekly for 10 weeks in the same cohort of mice. **c**, **d**, combined testes weight (**c**) and gonadosomatic index (combined testes weight/body weight; **d**). **e**, Representative stage VII–VIII seminiferous tubules stained for PLZF, KIT, SCP3, PNA and SOX9: nuclei counterstained with DAPI. **f**–**j**, Quantification of PLZF^+^ (**f**), KIT^+^ (**g**), SCP3^+^ (**h**), PNA^+^ (**i**) and SOX9^+^ (**j**) cells, normalized to tubule circumference (**f**: 20 images per mouse; **g**–**j**: 10 images per mouse). **k**, Representative tubules showing CASP3^+^/TUNEL^+^ peripheral and CASP3^−^/TUNEL^+^ luminal events. **l**,**m**, Quantification of CASP3^+^/TUNEL^+^ and CASP3^−^/TUNEL^+^ cells in peripheral (**l**) and luminal compartments (**m**). **n**, qPCR of fatty-acid metabolism, GPX4 defence, NRF2-response and iron-homeostasis transcripts, relative to ActB. **o**, **p**, Representative immunoblots and quantification of 4HNE (**o**) and GPX4 (**p**), normalized to GAPDH, black triangle indicates quantified band. **q**–**t**, Representative immunoblots (**q**) and quantification of FTH1 (**r**), FTL (**s**) and NCOA4 (**t**). **u**, FISH in RSs with X (red) or Y (green) signals and Y/X ratio (control: *n* = 6,527±135 Y-RSs vs *n* = 5,950±156 X-RSs; Y/X ratio mean 1.10; VE(+): *n* = 5,772±156 Y-RSs vs *n* = 5,702±149 X-RSs; mean Y/X ratio 1.01). Scale bars: **e**, 50 µm (overview) and **5** µm (inset); **k**, 50 µm; **u**, 5 µm. Data are mean ± s.d.; dots represent mice except for **b** where it represents the mean of each group. Statistics: two-way repeated measures-ANOVA with Geisser–Greenhouse correction (**b**); two-tailed Student’s *t*-test (**c**, **f**, **g**, **i**, **l**, **m**, **n**, **p**, **r**, **s**, **u**); Welch’s *t*-test (**o**); Mann–Whitney *U* test (**d**, **h**, **j**, **t**), exact *P* values are given. Sample sizes: **b**–**d**, *N* = 9 per group; all other panels, *N* = 6 per group.

We next assessed the transcriptional changes in RSs from aged mice. In VE(+) RSs, *Alox15* mRNA was significantly reduced, whereas expression of other lipid/oxidized lipid-metabolic genes including, *Gpx4*, NRF2-target genes and iron-related transcripts were unchanged relative to the aged controls (Fig. 5n). Nonetheless, 4HNE-modified protein levels were significantly reduced in VE(+) RSs (Fig. 5o). Among iron-handling proteins, in RSs from VE(+) mice, FTH1 and FTL protein levels were significantly reduced, whereas NCOA4 was increased (Fig. 5q–t), a pattern consistent with enhanced ferritin turnover, suggesting mobilization of ferritin-bound iron; however, whether this increases the labile iron pool or is coupled to iron utilization/export remains to be determined.

VE(+) also exerted an opposite effect on the RS sex chromosome composition compared to VE(−). In aged control testes, Y-bearing RSs outnumbered X-bearing RSs, whereas, in contrast, VE(+) restored the Y- and X-bearing RSs ratio to a near 1:1 ratio (control: mean Y/X ratio 1.10 [*n* = 6,527±135 Y-RSs vs *n* = 5,950±156 X-RSs]; VE(+): Y/X ratio 1.01 [*n* = 5,772±156 Y-RSs vs *n* = 5,702±149 X-RSs]; Fig. 5u). Taken together, the vitamin E deficiency and supplementation experiments show that in vitamin E deficient conditions, higher lipid peroxidation is associated with enrichment of Y-bearing RSs, whereas vitamin E supplementation in aged mice lowers lipid peroxidation and restores the RS sex ratio toward ∼1:1. These findings are consistent with preferential vulnerability of X-bearing RSs under conditions of elevated lipid peroxidation.

## Discussion

Aging in mammals is accompanied by testicular atrophy and progressive decline in spermatogenic capacity^28,41,42^. Here, we identified post-meiotic RSs as a particularly age-sensitive stage in which lipid peroxidation increases, coinciding with a compartment-specific shift in death signatures and an altered haploid sex-chromosome output. Dietary modulation of vitamin E reversed these phenotypes, indicating that membrane redox buffering can influence spermatogenic efficiency and the relative abundance of Y- and X-bearing haploid cells.

Aging increased germ-cell death, but not uniformly across the seminiferous epithelium. Peripheral layers showed an apoptotic signature consistent with prior work ^43,44^, whereas the luminal RS layer accumulated predominantly non-apoptotic TUNEL^+^ events. This stage specificity may reflect intrinsic vulnerabilities of RSs, whose membranes are enriched in peroxidation-prone PUFAs^15,45^, potentially pushing RSs closer to a lipid-damage threshold. RS-enriched fractions indeed displayed characteristics indicative of enhanced lipid peroxidation and reduced antioxidant capacity. The convergence of lipidomic oxidation signatures, altered antioxidant and iron-handling pathways, and bidirectional modulation by vitamin E supports a death modality that shares features with ferroptosis^18,23,46^.

A striking consequence of this RS vulnerability is a shift in haploid sex-chromosome composition. We observed a skew toward Y-bearing RSs in aged testes, indicating that the sex-chromosome ratio can deviate from ∼1:1 and persists in mature sperm. Because the skew is already present at the RS stage, it is unlikely to arise solely from selective processes during epididymal transit, although upstream meiotic drive and downstream quality-control mechanisms could still change the final output^47,48^. Round spermatids share gene transcripts across cytoplasmic bridges^49^, however, incomplete sharing of gene products across intercellular bridges provides a plausible route for divergence between X- and Y-bearing RSs, and geno-informativity analyses have identified incompletely shared gene transcripts relevant to lipid peroxidation pathways^50^. Interestingly, independent *in vitro* studies report an X-skew among surviving mature sperm following acute oxidative stress^51^. Although the direction of this bias differs from our observed RS-stage Y-skew, the two observations are not directly comparable given differences in developmental stage, context, and stress regime. Nonetheless, our dietary vitamin E data indicate that membrane redox buffering can influence RS haploid output, shifting the Y/X ratio toward or away from parity, consistent with the broader principle that redox conditions can bias sex-chromosome representation at the gamete level^52^, potentially via differential vulnerability of X- and Y-bearing cells.

This study has several limitations. First, although the lipid peroxidation profile and vitamin E sensitivity in RSs support a lipid damage state, definitive pathway assignment *in vivo* is constrained by the essential role of GPX4 in spermatogenesis, which limits straightforward genetic testing of this axis. Establishing causality will therefore require complementary approaches, such as perturbation of more genetically tractable suppressor pathways. Second, the basis of the aging-associated Y/X skew remains unknown because RS-enriched fractions were not sex-resolved and upstream or downstream selection may also contribute. Finally, we did not test whether vitamin E modifies the Y/X ratio in offspring sex ratio in aged males, which will require controlled mating studies.

Overall, our findings position RSs as a key stage at which aging-associated lipid redox stress may bias haploid sex-chromosome ratios (Extended Data Fig. 6). Defining how lipid peroxidation, antioxidant capacity and Y-versus X-linked vulnerabilities are coordinated *in vivo* will be important for understanding how male reproductive aging shapes sex-chromosome transmission.

## Methods

### Animals

Male C57BL/6J mice were maintained in the Animal Experiment Facility at the Tohoku University Graduate School of Medicine. All animals were housed in standard cages in a temperature- and humidity-controlled room on a 12-hour light/dark cycle (light on at 8 am) and had access to food and water ad libitum. All experimental procedures were approved by the Ethics Committee for Animal Experiments in Tohoku University (#2023-MED03701), and the animals were treated according to the National Institutes of Health (NIH) guide for the care and use of laboratory animals. For age-comparison experiments, C57BL/6J mice were obtained from Charles River Laboratories and were bred and maintained in-house. Mice were sacrificed at 3 months old (Young) or 18 months old (Aged).

### Vitamin E feeding

A separate cohort of male 1-month and 16-month-old C57BL/6J mice was purchased from Jackson Laboratories (JAX, Japan). Vitamin E diets were custom-made and ordered from Funabashi Farm, Japan. 1-month-old animals were fed control (containing 135 mg/kg vitamin E) or vitamin E-deficient diet (VE(−), containing <0.1 mg/kg vitamin E), and 16-month-old animals were maintained on control or vitamin E-supplemented diet (VE(+), containing 500 mg/kg vitamin E). Detailed ingredients are listed in Extended Data Table 1. All mice were raised on these diets for 10 weeks (70 days), covering two full cycles of spermatogenesis (35 days)^7^, after which mice were sacrificed at 3- and 18-months of age, respectively.

### Testis collection

Animals were sacrificed by sedation with isoflurane (MSD Animal health) and cervical dislocation. Testes were isolated, separated from the surrounding fat tissue and epididymis and weighed together. Samples were then fixed in 4% paraformaldehyde in phosphate buffered saline (PBS, 4% PFA, Nacalai-Tesque) for 2 hours at 4°C, cut in half, and fixed for an additional 18 hours. Cryoprotection was executed as follows: the testes samples were immersed, sequentially, in 10% sucrose dissolved in PBS for 1 hour and 20% sucrose in PBS overnight, both at 4°C. After cryoprotection, samples were frozen in Tissue-Tek optimal cutting compound (OCT, Sakura Finetek) on a metal pedestal using liquid nitrogen and stored in -80°C until further use. Cryosections (10µm) were obtained using a cryostat (Leica CM3050 S) and used for histological analyses.

### Immunohistochemistry (IHC)

Testicular sections on slide glasses (Matsunami) were stained using a previously established method^53^ with antibodies targeting anti-PLZF (1:500, sc28319, Santa Cruz), anti-SCP3 (1:500, ab97672, Abcam), anti-KIT (1:500, D13A2, Cell Signalling Technologies), anti-SOX9 (1:900, 5535, EMD Millipore), and anti-GPX4 (1:1,000, ab125066, Abcam). Sections were washed using Tris-buffered saline containing 0.1% Tween20 (TBST). Antigen retrieval was performed (except for anti-Kit) for 10 minutes in 0.01 M citrate buffer (pH 6.0) at 95°C. Afterwards, slides were washed with TBST and blocked with 3% bovine serum albumin (BSA) and 0.3% Triton X-100 in PBS for 1 hour at room temperature (RT). The slides were then incubated with primary antibodies diluted in blocking solution overnight at 4°C, washed with TBST, and incubated for 1 hour at RT with 4′,6-diamidino-2-phenylindole (DAPI, 1:1,000, Sigma) as well as either a Cy3- or Alexa488-conjugated secondary antibody (1:500, Jackson ImmunoResearch) or peanut agglutinin (PNA-Cy5, 1:2,000, Vector Laboratories) when staining the acrosome for round spermatid identification. Sections were washed again with TBST and mounted with Prolong Glass Antifade Mountant (Thermofisher). All images were obtained using a confocal laser-scanning microscope (LSM800, Carl Zeiss) and processed in Fiji (ImageJ).

### Cell death detection

Apoptotic or non-apoptotic cell death was identified by double labelling for terminal deoxynucleotidyl transferase dUTP nick end labelling (TUNEL, Merck Millipore) and an antibody against active-caspase-3 (1:500, BD Biosciences). TUNEL labelling was performed first and according to the manufacturer’s instructions. In short, slides with testis sections were washed in PBS and subjected to antigen retrieval by microwaving for 5 minutes in 0.01 M citrate buffer. TdT mixture was applied and incubated at 37°C for 1 hour, followed by incubation with an anti-digoxigenin antibody conjugated with fluorescein at RT for 30 minutes. Next, active-caspase-3 IHC was performed as described above with a slight modification: instead of DAPI, Hoechst-44432 (1:5,000) was used. Using Fiji, male germ line cells in the peripheral region of the seminiferous tissue and round spermatids in the luminal region were independently quantified.

### Fluorescent *in situ* hybridization (FISH)

FISH was performed according to a previously established protocol^54^. Slides with frozen sections of the testis were subjected to antigen retrieval in Histo-VT One (Nacalai-Tesque) for 30 minutes at 90°C and denatured using 50% formamide/2× saline-sodium citrate buffer (SSC), followed by dehydration using 70% and 100% ethanol. A custom mixture of X (conjugated with Cy3) and Y (conjugated with FITC) chromosome probes (Chromosome Science Labo) was applied to the slides, covered with a hybridization cover (Hybrislip, Thermofisher) and incubated at 80°C for 10 minutes. Further hybridization occurred overnight in a humid chamber at 37°C. To enhance probe signals, IHC was performed. Samples were blocked using 5% Blocking One solution (Nacalai-Tesque) for 1 hour at RT. Primary antibodies (anti-FITC, 1:2,000, A889, Invitrogen, and anti-Cy3, 1:2,000, sc-166894, Santa Cruz) were diluted in 5% Blocking One solution and incubated for 1 hour at RT. After washing with 4× SSC, Cy3- or Alexa488-conjugated secondary antibodies (1:2,000, Jackson ImmunoResearch) together with Hoechst-44432 (1:5,000, Thermofisher) were applied and incubated for 1 hour at RT. Slides were mounted with Prolong Glass Antifade Mountant (Thermofisher). All images were obtained using a fluorescence microscope (Keyence BZ-X series) and processed in Fiji. RSs were identified according to nuclear morphology and location within the tubule. Hybridization efficiency was determined by microscopic evaluation and images were deemed appropriate for analysis when, in more than 95% of cells, a sex chromosome could be assigned.

### Chemicals

Butylated hydroxytoluene (BHT, Cat#W218405-1KG-K), ammonium acetate for LC-MS LiChropur™ (Cat#5.33004) and dichloromethane (Cat#270997) were from Sigma-Aldrich. EquiSPLASH™ (Cat#330731) and UltimateSPLASH® ONE Mass Spec Standard (Cat#330820) were from Avanti Polar Lipids Inc. Methyl-tert-butyl ether (MTBE, Cat#34875), ammonium formate (NH4HCO2, Cat#55674) and formic acid (Cat#94318) were from Honeywell. Water (Cat#455), methanol (Cat#1428), acetonitrile (Cat#2697) and 2-propanol (Cat#1178) were from CHEMSOLUTE®.

### Lipid extraction from whole testis for lipidomic analysis

Lipid extraction from testes was performed using the methyl-tert-butyl ether (MTBE) method^55^. In short, approximately 100 mg of tissue was weighed into Precellys Lysing Kit tubes (P000912-LYSK1-A, Bertin Technologies) and 1 mL methanol was added. Samples were kept at 4°C and homogenized using a Precellys® 24 homogenizer (5,500 rpm, 3 times for 30 seconds, with a pause of 20 seconds between runs). Next, 500 µL of homogenate were transferred to a 10 mL screw-top glass vial (Chromacol 10-SV, Thermo) and spiked with internal standards (50 µL EquiSPLASH™ diluted 1:100 and 50 µL UltimateSPLASH® ONE Mass Spec Standard diluted 1:10, Avanti). Methanol (500 µL) and MTBE (3,330 µL) were added, and samples were vortexed using an Ohaus Multi-Tube Vortex Mixer (2,500 rpm for 2 minutes, pulsed), followed by incubation at 4°C for 20 minutes. A second vortexing step was performed (2,500 rpm for 2 minutes, pulsed) and samples were incubated at 4°C for an additional 20 minutes. Ice-cold water (830 µL) was added, samples were vortexed (2,500 rpm for 3 minutes, pulsed), and then centrifuged at 526 × g for 10 minutes at 4°C. The upper phase was transferred to a new screw-top vial. To the lower phase, MTBE (1,290 µL), methanol (388 µL), and water (323 µL) were added for re-extraction, followed by vortexing (2,500 rpm for 3 minutes, pulsed) and centrifugation at 526 × g for 10 minutes at 4°C. Upper phases were combined and evaporated to dryness using a RapidVap (Labconco) with the following settings: 45% speed, 35°C, 250 mbar for 30 minutes; 45% speed, 35°C, 150 mbar for 20 minutes; and 45% speed, 40°C, 50 mbar for 50 minutes. Dried extracts were resuspended in 200 µL 2-propanol and transferred to an amber 300 µL fixed insert vial (Chromacol 03-FIRV(A), Thermo) for LC–MS/MS analysis. A pooled sample was prepared by combining 7.5 µL of each extract. All solvents were supplemented with 0.01% (w/v) butylated hydroxytoluene (BHT) and cooled on ice prior to lipid extraction.

### Lipidomic analysis

Lipidomics was performed using hydrophilic interaction chromatography (HILIC) coupled to triple quadrupole mass spectrometry. HILIC was carried out on a SCIEX Triple Quad™ 7500 system equipped with a Luna® NH2 column (100 × 2 mm; 3 µm, 100 Å, Phenomenex). Lipids were separated by gradient elution using solvent A (acetonitrile/dichloromethane, 93:7, v/v) and solvent B (acetonitrile/water, 50:50, v/v), both containing 5 mM ammonium acetate. Solvent B was adjusted to pH 8.2 with NH3 (15%). Separation was performed at 35°C using the following gradient: 0 to 2 minutes, 0% B isocratic (flow rate 0.2 mL/min); 2 to 11 minutes, 0% to 40% B (flow rate 0.5 mL/min); 11 to 11.5 minutes, 40% to 70% B (flow rate 0.5 mL/min); 11.5 to 12.5 minutes, 70% to 100% B (flow rate 0.5 mL/min); 12.5 to 15 minutes, 100% B isocratic (flow rate 0.5 mL/min); 15 to 15.1 minutes, 100% to 0% B, followed by re-equilibration at 0% B for 2.5 minutes. The triple quadrupole mass spectrometer was operated with the following settings: TEM 500°C, GS1 45, GS2 70, CUR 45, CAD 9, IS −3,500 V. Data were acquired in multiple reaction monitoring (MRM) mode using a published assay (SCIEX technical note RUO-MKT-02-8477-C). For quantification, the area under the curve for each precursor mass to fragment mass transition was integrated using SCIEX OS software and normalized to appropriate internal standards from UltimateSPLASH® ONE (Avanti), including PC(17:0_22:4(d5)), PE(17:0_18:1(d5)), LPC(17:0(d5)), and LPE(17:0(d5)). Isotopic correction was performed using LICAR^56^.

### Round spermatid (RS) isolation

Isolation was carried out according to a previously established protocol^57^. Briefly, Fresh testicular tissue was harvested from young or aged male mice followed by removal of the epididymis and tunica albuginea. A single cell suspension was obtained via serial digestion with Collagenase IV (1mg/ml, Sigma) and Trypsin (0.6 mg/ml, Worthington) / DNAse I (>3.2 kU/ml, Sigma) dissolved in Krebs bicarbonate buffer. The cell suspension was loaded on a 1-5% BSA / Krebs buffer gradient and allowed to sediment for 1.5 hour on ice at 1 unit gravity. Twenty-eight fractions of 1 mL were collected and analyzed for purity using PNA (1:2,000, Rhodamine, Vector Laboratories) and DAPI (1:1,000, Sigma). Only fractions #4-7 were routinely enriched with RSs (>80%, Extended Data Fig. 3). Therefore, in subsequent isolations, fractions #4-7 were collected and pooled. The fractions were washed with Krebs buffer to remove excess BSA and immediately used for RNA and protein isolation.

### Simultaneous RNA and Protein isolation

RNA and protein were simultaneously isolated using the AllPrep DNA/RNA/Protein mini kit (Qiagen) according to the manufacturer’s protocol, with one minor modification: the protein fraction was air dried and then vortexed for 1 minute in SDS buffer (containing 4% SDS, 20% sucrose, 10 mM Tris-HCl pH 6.8 and 1× ethylenediaminetetraacetic acid (EDTA)-free protease inhibitor (Roche), followed by incubation at 37°C for 30 minutes. The protein sample was vortexed and pelleted via centrifugation. Supernatant was transferred to a new tube and protein concentration was determined using the bicinchoninic acid assay (BCA, Nacalai-Tesque). RNA was assessed for concentration and purity using the Nanodrop (Thermofisher). Isolated RNA and protein were stored at -80°C until further use.

### cDNA synthesis and quantitative polymerase chain reaction (qPCR)

RNA isolated from pooled round spermatid fractions (1 µg) was reverse transcribed into cDNA using the SuperScript III™ First-Strand Synthesis System for RT-PCR (Invitrogen). cDNA (2 µl, 1:10 diluted) was subsequently used for qPCR using Taq Pro Universal SYBR qPCR Master Mix (Vazyme). Primer sequences used in this study can be found in Extended Data Table 2. Assay specificity was verified by single-peak melt curves and a single band of the expected size on agarose gel. For *Gpx4* transcript analysis, three qPCR assays were used^39^: a primer set specific for the *nGpx4* transcript, a primer set specific for the mitochondrial-targeted *Gpx4* transcript (*mGpx4*), and a primer set amplifying a region shared by mitochondrial and cytosolic transcripts (*mGpx4+cGpx4*). Ct values were normalized to *ActB* (ΔCt). To estimate the cytosolic component, expression values were converted to linear scale (2^−ΔCt^), and cytosolic expression was calculated as ***E***_*c*_ = ***E***_*m*+*c*_ − *E*_*m*_, where ***E***_*m*+*c*_ is the signal from the assay amplifying the shared mitochondrial+cytosolic transcripts and *E*_*m*_is the mitochondrial-specific assay. Group comparisons were performed on log2-transformed expression values.

### Western blot

Protein samples (5 µg) were separated by SDS-PAGE on 12% TGX™ FastCast™ acrylamide gels (Bio-Rad) at 150 V for 50 minutes and transferred to 0.2 µm polyvinylidene difluoride (PVDF) membranes (Trans-Blot Turbo Mini PVDF Transfer Packs; Bio-Rad) using the Trans-Blot Turbo system (2.5 A, 25 V, 3 minutes). Membranes were blocked in 10% Intercept TBS blocking buffer (Licor; GPX4, ActB), 5% Blocking One (Nacalai-Tesque; 4HNE, GAPDH), or 5% skimmed milk in TBST (FTH1, FTL and NCOA4) and incubated overnight at 4°C with primary antibodies against GPX4 (1:1,000, ab125066, Abcam), 4HNE, (1:1,000, ab46545, Abcam), ACTB (1:2,500, sc-47778, Santa Cruz), GAPDH (1:4,000, ab8245, Abcam), FTH1 (1:1,000, ab183781, abcam) FTL (1:1,000, ab69090, abcam) and NCOA4 (1:1,000, ab314553, abcam). After washing with Tris-buffered saline containing 0.1% Tween-20 (TBST), membranes were incubated for 1 hour at room temperature with HRP-conjugated secondary antibodies: for GPX4 and ACTB, W4011 (1:10,000) and W4021 (1:5,000) in 10% Intercept; for 4HNE and GAPDH, both 1:10,000 in 5% Blocking One; and for FTH1, FTL and NCOA4, both 1:10,000 in ‘Can Get Signal?’ (Toyobo). Membranes were washed again in TBST, and signals were visualized using a ChemiDoc Touch Imaging System (Bio-Rad). Band intensities were quantified in Fiji and normalized to ACTB or GAPDH as specified.

### Flow cytometric analysis of mature sperm

The caudal epididymis was excised from two testes, incised 4 times, and transferred to a dish containing 2 mL prewarmed (37°C) phosphate buffered saline (PBS) with 2% FBS. Viable mature sperm were allowed to swim out into the medium for 10 minutes in a 37°C, 5.0% CO_2_ incubator. The sperm suspension was filtered through a 40 µm PET cell strainer and stained with Hoechst (5 µg/mL) for 45 minutes at 35°C in the dark, mixing every 15 minutes. Samples were kept on ice until analysis, which proceeded within 1 hour. Directly before analysis, 2 mL cell suspensions were filtered through a 35 µm strainer cap attached to a 5 mL Polystyrene Round Bottom FACS tube (Falcon). Cells were subjected to flow cytometry using an LSRFortessa cell analyzer (BD Biosciences). Hoechst was excited using the 355 nm UV laser and detected using a 450/50 bandpass filter. Forward Scatter (FSC) and Side Scatter (SSC) were detected using a 488 nm laser and a 488/10 bandpass filter. Flow rate was set to achieve 2,500-3,500 events/second. A minimum of 100,000 events within a specified gate (Fig 3d) were recorded using the BD FACSDiva Software (BD Biosciences). Data analysis was done using FlowJo software v10.10.0 (BD Life Sciences).

### Statistical analysis

Statistical information for individual experiments is shown in the corresponding figure legends. Values are presented as mean ± standard deviation. Data were first analysed for normality using the Shapiro–Wilk test. Depending on distribution and variance, statistical comparisons between groups were performed using Student’s *t*-test, Welch’s *t*-test, or the Mann–Whitney *U* test. Longitudinal effects were analysed using two-way ANOVA (fixed effects: diet, week and bodyweight; random effect: individual mouse, repeated measures). Results were considered significant at *P* < 0.05. Statistical analyses were conducted using GraphPad Prism 10 (GraphPad Software).

## Acknowledgements

We thank the Japanese Society for the Promotion of Science KAKENHI grant number #17K19372, #22K19322, and #25K22536, The Canon Foundation awarded to N.O., the establishment of university fellowships towards the creation of science technology innovation, grant number JPMJFS2102, JPMJSP2114 awarded to J.G., and JST FOREST Program (JPMJFR2213) awarded to E.M. for financial support. M.C. received funding from the European Research Council (ERC) under the European Union’s Horizon 2020 research and innovation program (grant agreement no. GA 884754). We thank Sayaka Makino and Asuka Takehara for animal care and all other members of the Developmental Neuroscience laboratory at Tohoku University for their useful comments. We also thank Prof. Junken Aoki (Graduate School of Pharmaceutical Sciences, The University of Tokyo) for advice on lipidomic analysis and Dr. Ibuki Kusumoto (Laboratory of Food Function Analysis, Tohoku University) for technical support.

## Author contributions

Study design: J.G., T.K., W.G., E.M., N.O.; Data collection: J.G., T.K., J.I., L.G., B.H.; Data analysis: J.G.; J.I., B.H., M.A.; Drafted the manuscript: J.G.; Edited the manuscript: T.K., W.G., E.M., M.C., N.O.; Funding acquisition: J.G., E.M., M.C., N.O.

---

**Extended data Fig. 1:**
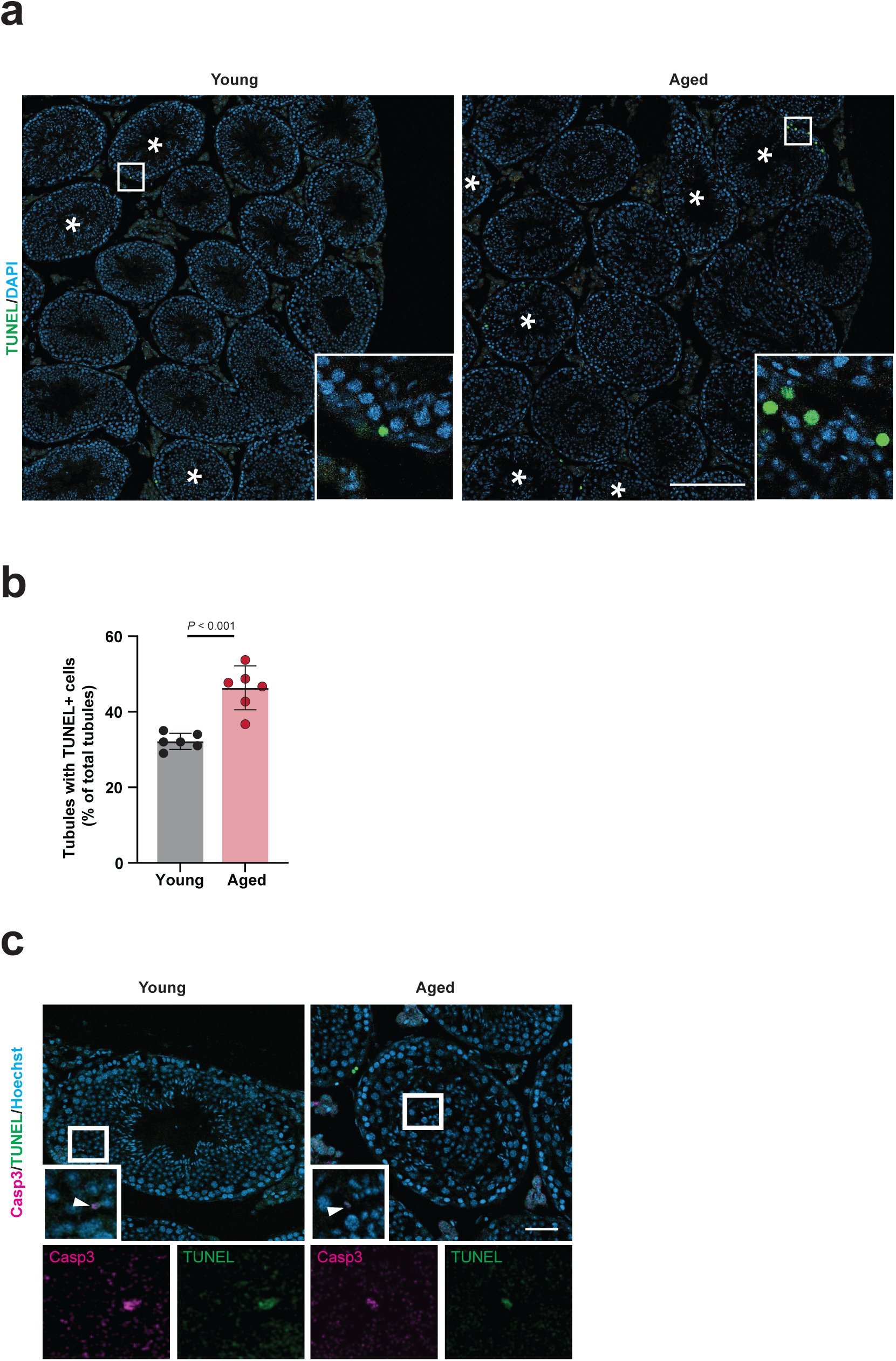
Age-associated cell death is increased in seminiferous tubules. **a**, Representative images of tubules containing TUNEL^+^ cells in young and aged mice. Asterisks indicate tubules containing TUNEL^+^ cells. Scale bar: 200 µm. **b**, Percentage of seminiferous tubules containing cells TUNEL^+^ cells relative to the total number of tubules within a section. **c**, Representative images of CASP3^+^/TUNEL^+^ round spermatids in young and aged testis sections. Scale bar: 50µm. Data is presented as mean ± s.d. Significance was determined by two-tailed Student’s *t*-test (**b**). Exact *P* value is given. Sample size: *N* = 6 in each group. Each dot represents one mouse.

**Extended data Fig. 2:**
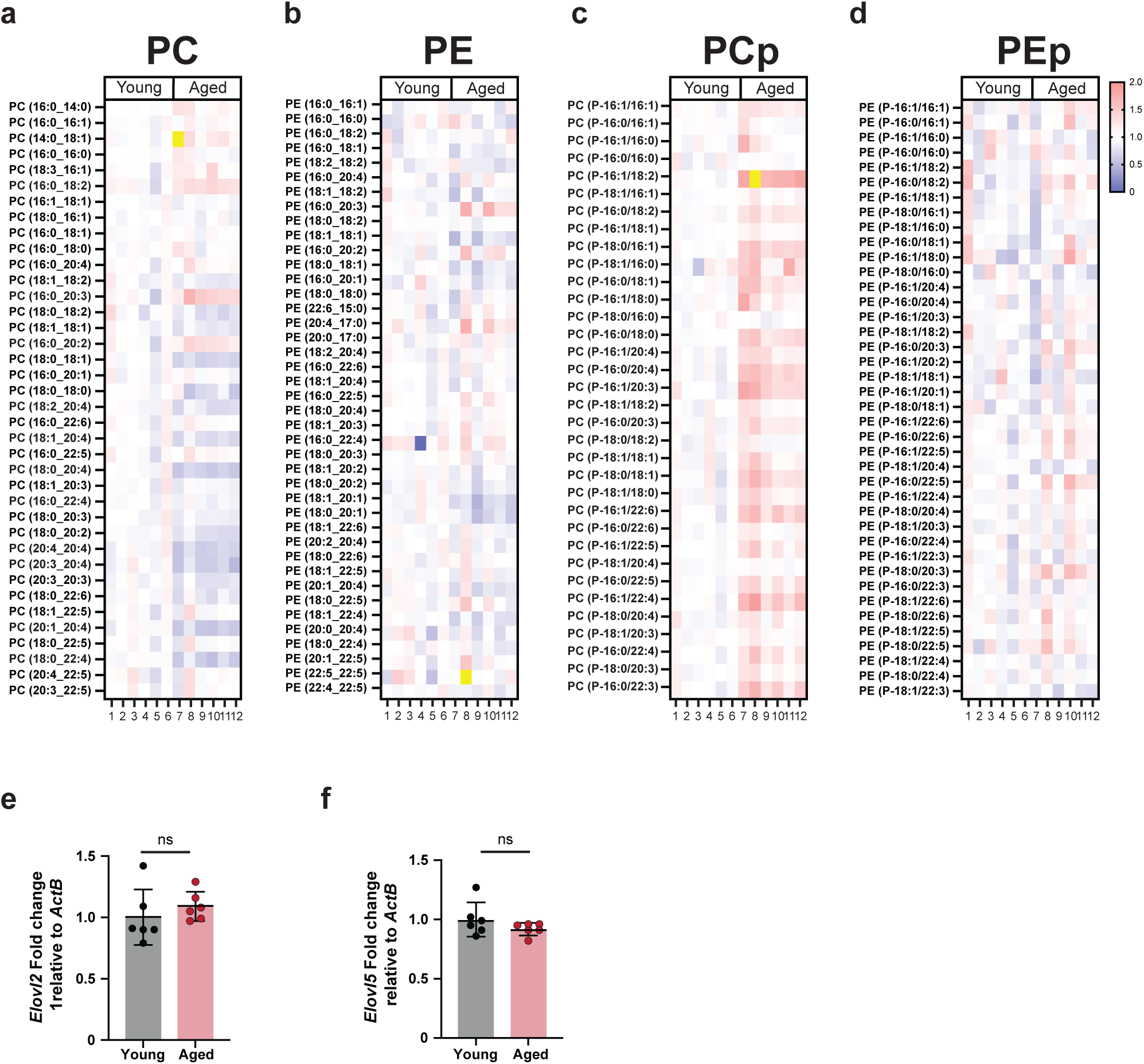
Aging shifts phosphatidylcholine composition toward plasmalogen PC. **a**-**d**, Heatmaps showing relative abundances of individual phospholipid species in young and aged whole testis: diacyl phosphatidylcholine (PC; **a**), diacyl phosphatidylethanolamine (PE; **b**), plasmalogen phosphatidylcholine (PCp; **c**) and plasmalogen phosphatidylethanolamine (PEp; **d**). Columns represent biological replicates and rows represent lipid species. Values are scaled to show relative differences across samples. **e**, **f**, mRNA level quantification analysis of *Elovl2* (**e**) and *Elovl5* (**f**) expression in young and aged round spermatids. Data is presented as mean ± s.d. Significance was determined by two-tailed Student’s *t*-test (**e, f**). Sample size: *N* = 6 in each group. Each dot represents one mouse.

**Extended data Fig. 3:**
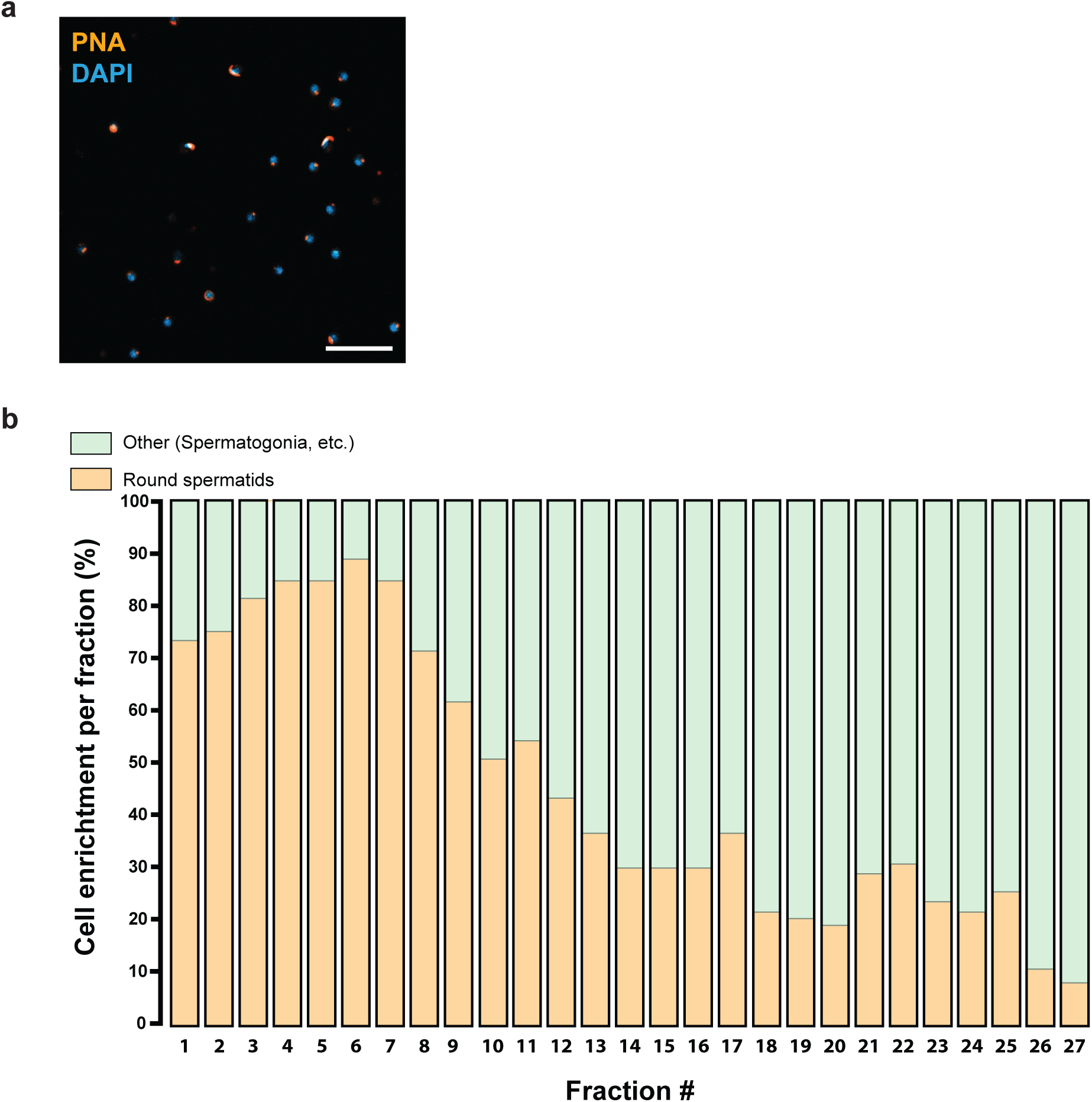
Enrichment profile of round spermatids across fractions. a, Representative image of the enriched cell suspension (fraction #3) stained with peanut agglutinin (PNA) to label the acrosome and DAPI to label nuclei. Scale bar: 20µm. **b,** Representative cellular composition of fractions showing the percentage of round spermatids versus other germ-cell populations (including spermatogonia and other cells). Each bar represents one fraction; stacked colors indicate the relative contribution of round spermatids (orange) and other cells (green).

**Extended data Fig. 4:**
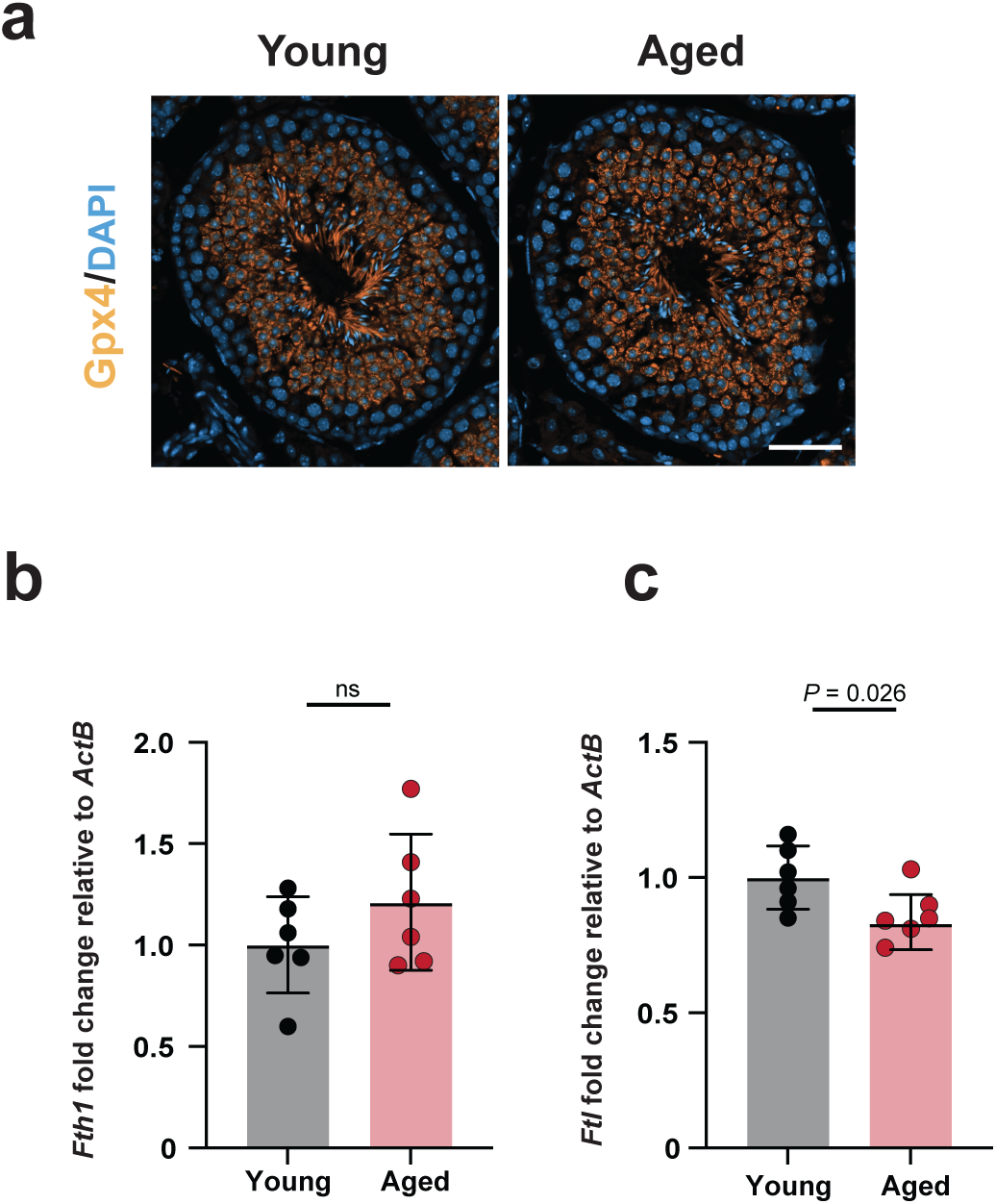
GPX4 localization and ferritin expression in testicular aging a, Representative images of GPX4 localization in seminiferous tubules from young and aged testes, with DAPI counterstain. Scale bar: 50µm. **b, c,** mRNA level quantification of iron-handling transcripts in round spermatids from young and aged mice: *Fth1* (**b**) and *Ftl* (**c**), relative to *ActB*. Data are presented as mean ± s.d. Significance was determined by two-tailed Student’s *t*-test (**b, c**). Exact *P* values are given. Sample size: *N* = 6 in each group. Each dot represents one mouse.

**Extended data Fig. 5:**
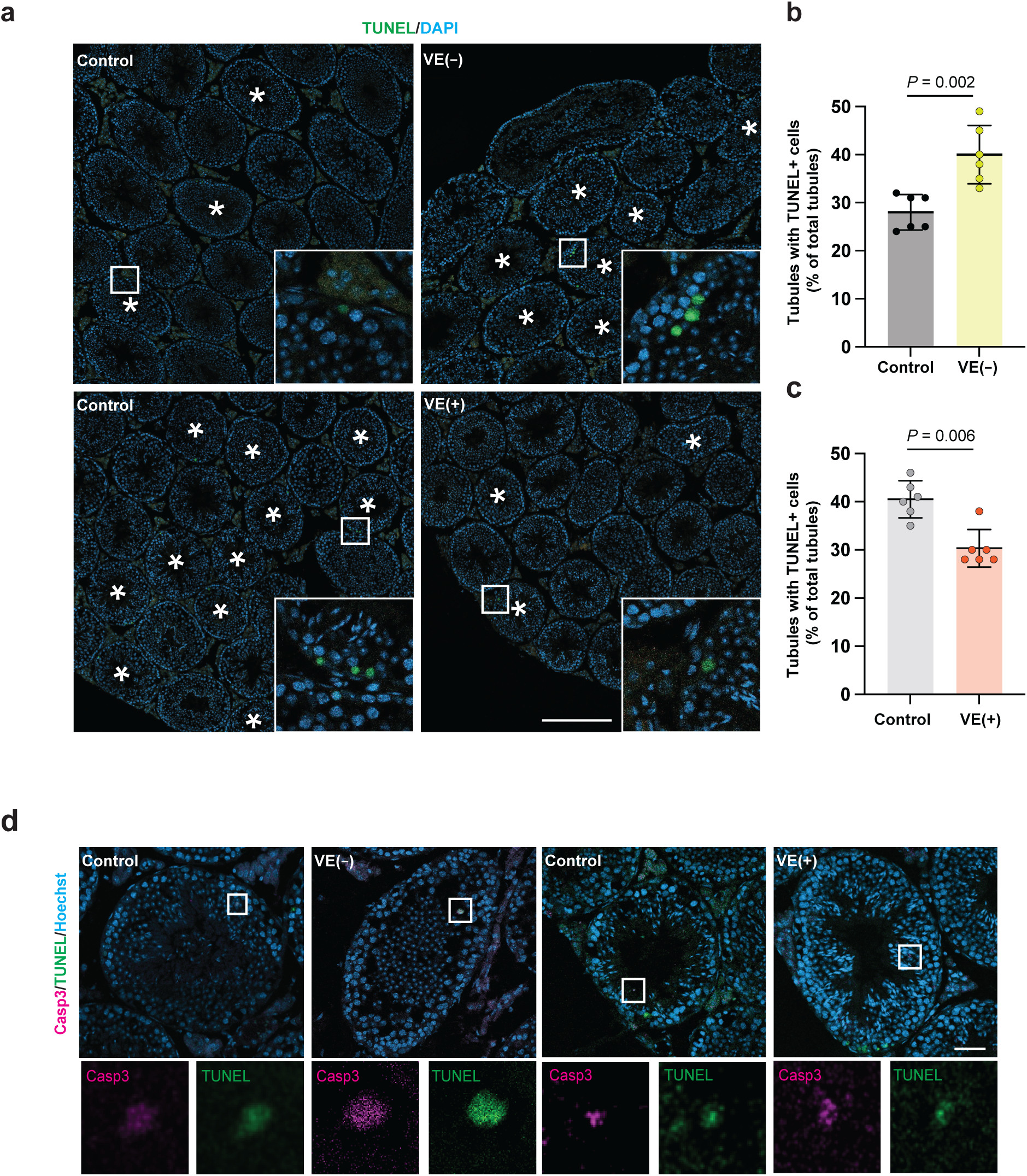
Dietary vitamin E modulates cell death in young and aged testes a, Representative TUNEL staining in testis sections from young control (3-months), vitamin E–deficient (VE(−)), aged (18-months) control and aged vitamin E–supplemented (VE(+)) mice; asterisks mark tubules containing TUNEL^+^ cells. Scale bar: 200 µm. **b, c,** Quantification of the percentage of tubules containing TUNEL^+^ cells in young mice (control vs VE(−); **b**) and aged mice (control vs VE(+); **c**). **d,** Representative images of CASP3^+^/TUNEL^+^ round spermatids in young and aged testis sections. Scale bars: 50µm. Data are presented as mean ± s.d. Significance was determined by two-tailed Student’s *t*-test (**b, c**). Exact *P* values are given. Sample size: *N* = 6 in each group. Each dot represents one mouse.

**Extended Data Fig. 6:**
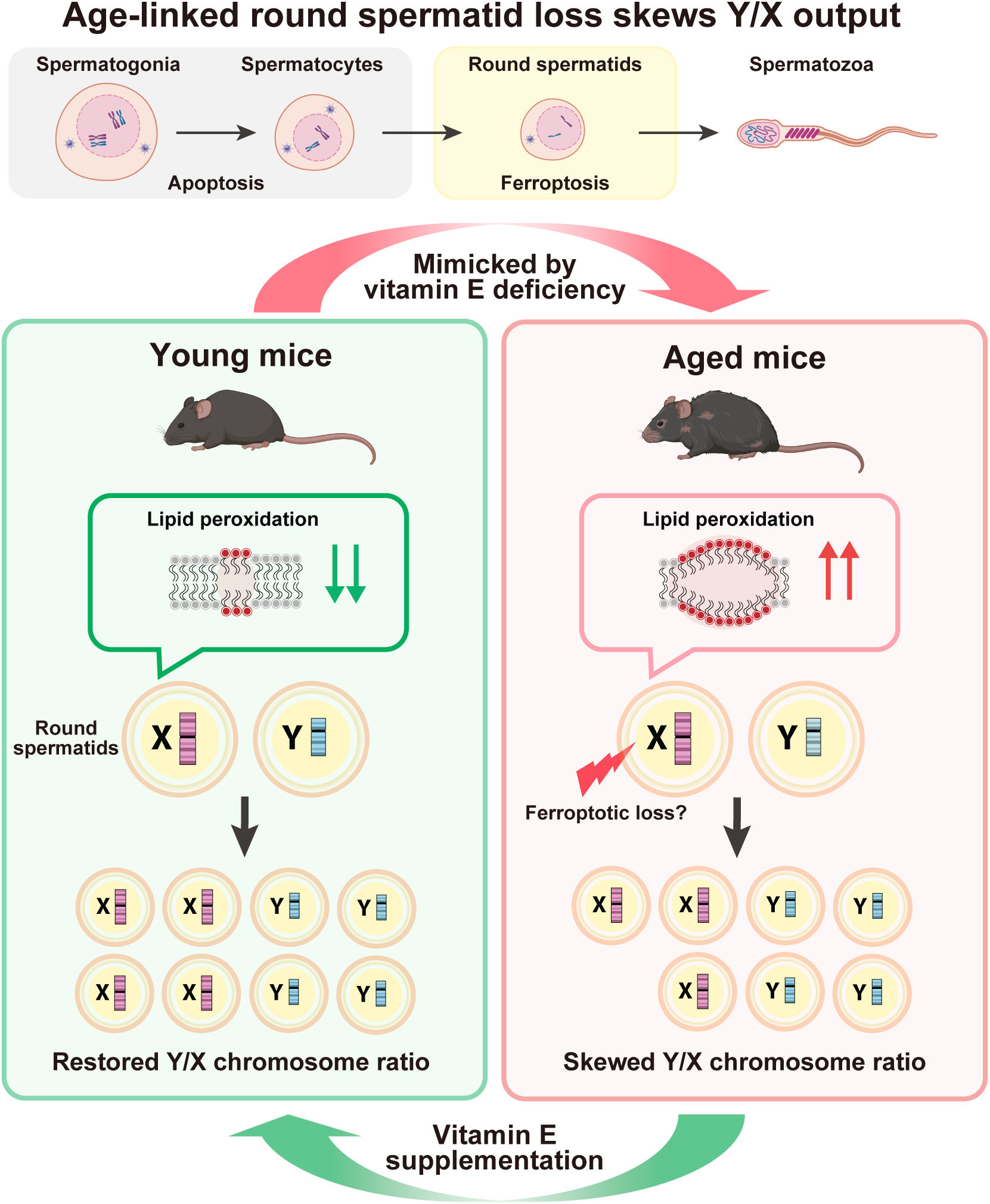
Age-linked round spermatid loss skews Y/X output. Graphical summary of the model supported by this study, linking aging-associated lipid redox stress to altered haploid sex-chromosome output. Along the male germline lineage, cell death is enriched in pre-meiotic and meiotic stages, whereas post-meiotic round spermatids (RSs) exhibit heightened lipid peroxidation and ferroptosis-like vulnerability. In young mice, lipid peroxidation is low and X- and Y-bearing RSs are maintained, resulting in a balanced Y/X output. In aged mice, lipid peroxidation is increased and RS loss is associated with a skewed Y/X chromosome ratio. Vitamin E deficiency phenocopies the aging-associated increase in lipid peroxidation and RS, whereas vitamin E supplementation reduces lipid peroxidation and restores Y/X balance. Arrows denote the direction of change; the lightning symbol indicates RS loss. Illustration was created in BioRender. Germeraad, J. (2026) https://BioRender.com/k04h4kg

