## Supplementary tables for "Ferroptotic vulnerability of post-meiotic germ cells skews sex chromosome ratios in aged testis"

**Diets based on AIN76 feed by Funabashi farm**

| Control diet (Per kg) | |
|---|---|
| **Vitamins** | **Quantity** |
| Vitamin A (IU) | 4000 |
| Vitamin $D_3$ (IU) | 1000 |
| **Vitamin E (α, β, γ, δ) (mg)** | **135** |
| Vitamin $K_3$ (μg) | 50 |
| Vitamin $B_1$ (mg) | 6 |
| Vitamin $B_2$ (mg) | 6 |
| Vitamin $B_6$ (mg) | 7 |
| Vitamin $B_{12}$ (μg) | 10 |
| Biotin (μg) | 200 |
| Folic acid (mg) | 2 |
| Calcium pantothenate (mg) | 16 |
| Nicotinic acid (mg) | 30 |
| Choline bitartrate (g) | 2 |
| **Minerals** | **Quantity** |
| Calcium (mg) | 5000 |
| Phosphorus (mg) | 4000 |
| Magnesium (mg) | 500 |
| Sodium (mg) | 1000 |
| Potassium (mg) | 3600 |
| Iron (mg) | 35 |
| Copper (mg) | 6 |
| Zinc (mg) | 30 |
| Mangan (mg) | 54 |
| Selenium (mg) | 0 1 |
| Chromium (mg) | 2 |
| Iodine (mg) | 0 2 |
| Chlorine (mg) | 1560 |
| Sulfate radical (mg) | 1000 |

| Vitamin E deficient diet (Per kg) | |
|---|---|
| **Vitamins** | **Quantity** |
| Vitamin A (IU) | 4000 |
| Vitamin $D_3$ (IU) | 1000 |
| **Vitamin E (α, β, γ, δ) (mg)** | **<0.1** |
| Vitamin $K_3$ (μg) | 50 |
| Vitamin $B_1$ (mg) | 6 |
| Vitamin $B_2$ (mg) | 6 |
| Vitamin $B_6$ (mg) | 7 |
| Vitamin $B_{12}$ (μg) | 10 |
| Biotin (μg) | 200 |
| Folic acid (mg) | 2 |
| Calcium pantothenate (mg) | 16 |
| Nicotinic acid (mg) | 30 |
| Choline bitartrate (g) | 2 |
| **Minerals** | **Quantity** |
| Calcium (mg) | 5000 |
| Phosphorus (mg) | 4000 |
| Magnesium (mg) | 500 |
| Sodium (mg) | 1000 |
| Potassium (mg) | 3600 |
| Iron (mg) | 35 |
| Copper (mg) | 6 |
| Zinc (mg) | 30 |
| Mangan (mg) | 54 |
| Selenium (mg) | 0 1 |
| Chromium (mg) | 2 |
| Iodine (mg) | 0 2 |
| Chlorine (mg) | 1560 |
| Sulfate radical (mg) | 1000 |

| Vitamin E supplemented diet (Per kg) | |
|---|---|
| **Vitamins** | **Quantity** |
| Vitamin A (IU) | 4000 |
| Vitamin $D_3$ (IU) | 1000 |
| **Vitamin E (α, β, γ, δ) (mg)** | **500** |
| Vitamin $K_3$ (μg) | 50 |
| Vitamin $B_1$ (mg) | 6 |
| Vitamin $B_2$ (mg) | 6 |
| Vitamin $B_6$ (mg) | 7 |
| Vitamin $B_{12}$ (μg) | 10 |
| Biotin (μg) | 200 |
| Folic acid (mg) | 2 |
| Calcium pantothenate (mg) | 16 |
| Nicotinic acid (mg) | 30 |
| Choline bitartrate (g) | 2 |
| **Minerals** | **Quantity** |
| Calcium (mg) | 5000 |
| Phosphorus (mg) | 4000 |
| Magnesium (mg) | 500 |
| Sodium (mg) | 1000 |
| Potassium (mg) | 3600 |
| Iron (mg) | 35 |
| Copper (mg) | 6 |
| Zinc (mg) | 30 |
| Mangan (mg) | 54 |
| Selenium (mg) | 0 1 |
| Chromium (mg) | 2 |
| Iodine (mg) | 0 2 |
| Chlorine (mg) | 1560 |
| Sulfate radical (mg) | 1000 |

| Composition of refined feed (Per kg) | (%) |
|---|---|
| Moisture | 9 |
| Crude protein | 16 6 |
| Crude fat | 4 9 |
| Crude fiber | 2 9 |
| Crude ash content | 2 2 |
| Carbohydrates | 64 4 |
| Calories | 3690 kcal |

**Primer sequences**

| Primer name | Sequence 5' -> 3' |
|---|---|
| Acsl4_FW | CCAAAGAACACCATTGCCATT |
| Acsl4_RV | AAGTCTGTGCTGCAATCATCCA |
| Alox15_FW | GTACGCGGGCTCCAACAACGA |
| Alox15_RV | TCTCCGGGGCCCTTCACAGAA |
| mGpx4_FW | GAGATGAGCTGGGGCCGTCTGA |
| mGpx4_RV | ACGCAGCCGTTCTTATCAATGAGAA |
| nGpx4_FW | AGTTCCTGGGCTTGTGTGCATCC |
| nGpx4_RV | ACGCAGCCGTTCTTATCAATGAGAA |
| cmGpx4_FW | CGCCTGGTCTGGCAGGCACCA |
| cmGpx4_RV | ACGCAGCCGTTCTTATCAATGAGAA |
| ActB_FW | TATGCCAACACAGTGCTGTC |
| ActB_RV | ACCGATCCACACAGAGTACTTG |
| Sod1_FW | CAGAAGGCAAGCGGTGAAC |
| Sod1_RV | CAGCCTTGTGTATTGTCCCCATA |
| Catalase_FW | GGACGCTCAGCTTTTCATTC |
| Catalase_RV | TTGTCCAGAAGAGCCTGGAT |
| Nrf2_FW | CAGTGCTCCTATGCGTGAA |
| Nrf2_RV | GCGGCTTGAATGTTTGTC |
| Hmox1_FW | GGAAATCATCCCTTGCACGC |
| Hmox1_RV | TGTTTGAACTTGGTGGGGCT |
| Acsl6_FW | AAGCAGTCGGAAGAAGTGGAG |
| Acsl6_RV | GGCATCGTCATAGTAATGGGTAAG |
| Elovl2_FW | TCAATGCTTTCTTGGACAACATG |
| Elovl2_RV | GGTAAGAGTCCAGCAGGAACCA |
| Elovl5_FW | ATGGACACCTTTTTCTTCATCCTT |
| Elovl5_RV | ATGGTAGCGTGGTGGTAGACATG |
| Fth1_FW | GGCAAAGTTCTTCAGAGCCA |
| Fth1_RV | CATCAACCGCCAGATCAAC |
| Tfr1_FW | TCAAGCCAGATCAGCAATTCTC |
| Tfr1_RV | AGCCAGTTTCATCTCCACATG |
| Ftl1_FW | CAGCCTGGTCAATTTGTACCT |
| Ftl1_RV | GCCAATTCGCGGAAGAAGTG |
